# Evolutionary instability drives structural diversity and disease susceptibility at the 16p12.2 locus

**DOI:** 10.64898/2026.03.04.709583

**Authors:** Corrine Smolen, Santhosh Girirajan

## Abstract

Extensive duplication in the African great ape lineage has led to substantial instability of chromosome 16p. We examined the sequence structure and evolutionary history of the 16p12.2 locus in 570 diverse human haplotypes and seven non-human primates. Human haplotypes vary greatly in size and exhibit ancestry-biased structure. We identify 5-14 clusters of distinct architecture at three segmental duplication (SD) blocks, generating 21 unique haplotype configurations. Two duplicons within these SDs, D5 and D6, mediate the neurodevelopmental disorder-associated 16p12.1 deletion; however, exact breakpoint positions and local sequence architecture vary across families. The region has toggled between orientations over the past 25 million years, and we identify 32 inversions in humans mediated by distinct duplicons. Evolutionary analyses reveal incomplete lineage sorting, interlocus gene conversion, and lineage-specific expansions, including human-specific expansions of D5 and D6. These findings highlight the evolutionary instability at 16p12.2 driving structural diversity and deletion susceptibility in humans.

---

Initial analysis of the human genome revealed that highly repetitive low copy repeats, termed segmental duplications (SDs), are non-randomly distributed and enriched at over 100 hotspots of non-allelic homologous recombination (NAHR), extending beyond classical loci such as 22q11.2 and 17p11.2^1–5^. Comprising approximately 7% of the human genome^6^, SDs are more abundant in humans than other mammals^7^. This pattern can be traced to a burst of duplication events in the common ancestor of African great apes^8^. In particular, the short arm of human chromosome 16 has undergone rapid expansion of duplicons, a sequence subunit shared across multiple SD blocks, by frequent intrachromosomal transposition events along the great ape and human lineages^9,10^. The high sequence identity of duplicons on chr16p has created substrates for recurrent NAHR, contributing to deletions and duplications at 16p11.2^11^, 16p12.2^12^, and 16p13.11^13^, regions associated with several neurodevelopmental disorders, including developmental delays, autism^14,15^, schizophrenia^16,17^, and epilepsy^18^. For instance, Nuttle and colleagues identified a 95 kbp duplicon containing *BOLA2* that expanded specifically in the human lineage leading to increased genomic instability and susceptibility to recurrent 16p11.2 rearrangements^19^.

Another locus on chr16p, 16p12.2, is a complex region containing three large SD blocks, BP1-BP3 (**Fig. 1A**). Highlighting its complexity, the region was misaligned in past reference genomes^20^ and has only been corrected in the recent telomere to telomere CHM13 genome^21^. This region is the locus for the 16p12.1 deletion (on hg18 when originally described, maps to 16p12.2 on more recent reference genomes), an ∼520 kbp deletion which encompasses eight genes and occurs in approximately one in 1,400-2,000 individuals^22^. It was originally described in children with developmental delays^12^ but has since been associated with a wide range of clinical phenotypes at varying levels of severity and penetrance^22–26^. Previous studies have sought to understand the structure of the locus^20,27^, however, the complete architecture of SDs mediating the 16p12.1 deletion, exact deletion breakpoints, and the evolutionary history of this region across primate lineages are not completely understood.

**Figure 1.**
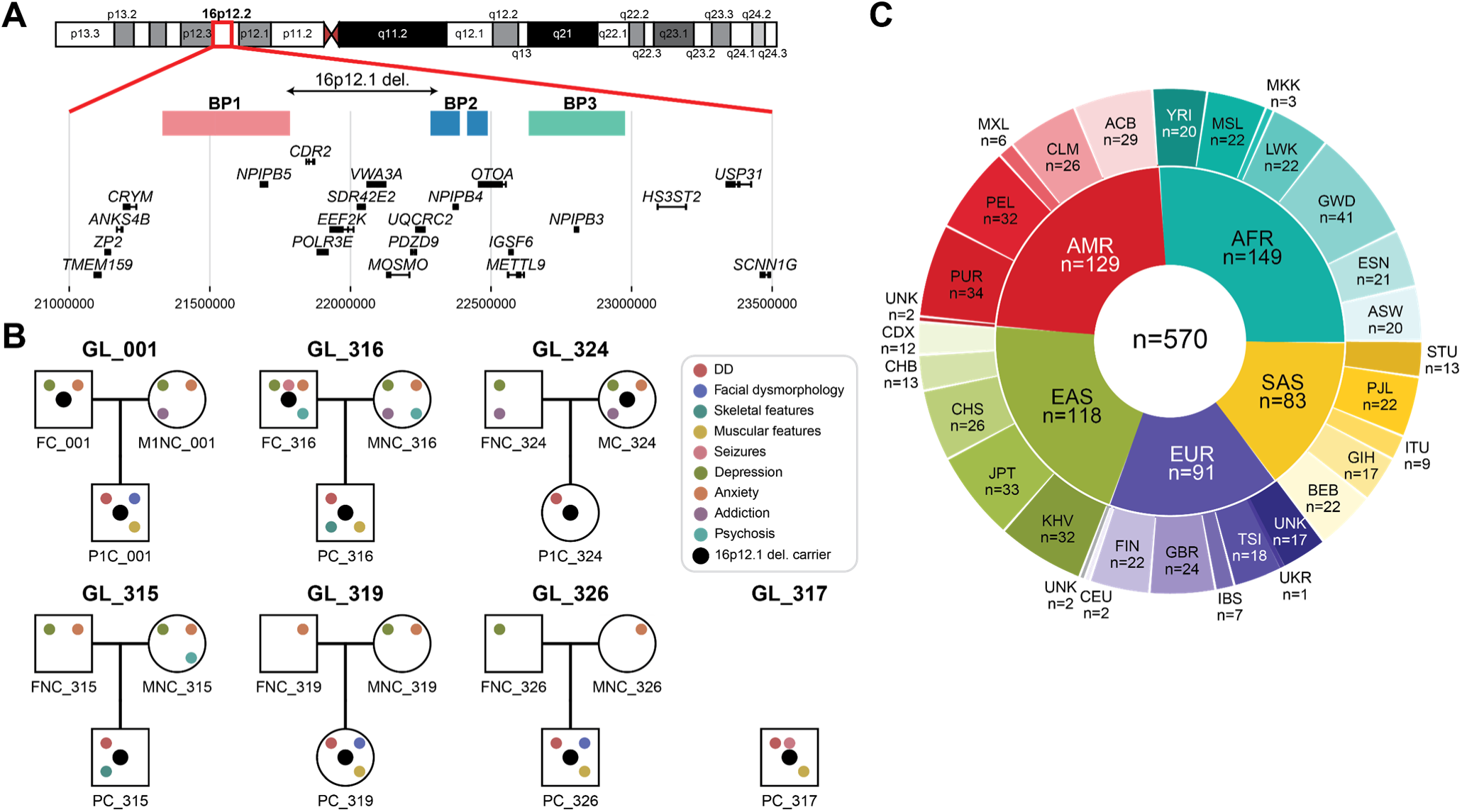
16p12.2 locus and cohort. **(A)** 16p12.2 locus on the CHM13-T2T reference genome. **(B)** 16p12.1 deletion families with variable phenotypes. **(C)** Population descriptors of haplotypes available for analysis of 16p12.2 region, including 21 haplotypes from 16p12.1 deletion families. Populations are labelled according to the population descriptors provided by HPRC and HGSVC, when available, or through descriptors assigned to 16p12.1 deletion families based on self-reported ancestry information. UKR describes Ukrainian participants. UNK describes participants where the superpopulation to which they belong is known (i.e. European, African), but specific population is unknown.

Previous studies using microarrays^28^, fluorescence *in situ* hybridization^4^, optical mapping^29,30^, and short read whole genome sequencing^31^ lacked the resolution^32–35^ required to fully characterize SDs and associated recurrent rearrangements. Due to these limitations, the architecture of CNVs and the surrounding regions have only recently begun to be fully described for a subset of recurrent CNVs^36,37^. This can be attributed to advancement in sequencing technologies^38–40^ that allowed for the increased availability of high quality genome assemblies from diverse individuals^41^ and more complete reference genomes^42^, providing avenues for investigation of previously inaccessible parts of the genome.

Here, we performed long read whole genome sequencing from seven families with the 16p12.1 deletion and generated haplotype-phased assemblies. We additionally leveraged more than 500 phased genomes from the Human Pangenome Reference Consortium (HPRC)^43^ and the Human Genome Structural Variation Consortium (HGSVC)^44^ to define 21 unique SD architecture configurations and identified duplicons in the region. We further examined recombination events in the deleted allele from a proband with a *de novo* deletion to identify potential NAHR events and localized variable deletion breakpoints across families to two duplicon units present in BP1 and BP2. We also assessed the evolutionary history of the locus, and identified evolutionary “toggling” of the regions flanked by SDs, using recent high quality reference genomes from six ape species^45^ and rhesus macaque^46^. We additionally found 32 inversions in the region mediated by multiple distinct duplicons. We finally tracked the evolutionary history of individual duplicons and identify patterns of incomplete lineage sorting, lineage-specific expansions, and interlocus gene conversion. Together, these data reveal that the diverse architecture and deletion susceptibility in modern humans are driven by the evolutionary instability of the locus.

## RESULTS

### Identification and characterization of 16p12.2 region haplotypes

We focused our analysis on the 2.5 Mbp region encompassing the variably expressive 16p12.1 deletion region (CHM13 coordinates: chr16:21000000-23500000). This interval contains three segmental duplication (SD) blocks (BP1, BP2, and BP3) and 21 protein-coding genes (**Fig. 1A**). We generated 30X PacBio HiFi whole genome sequencing (WGS) data from six family trios and one singleton proband with the 16p12.1 deletion (**Fig. 1B**). These trios include three individuals with *de novo* and three probands with inherited deletions. We note that one of the *de novo* cases, PC_326, carried a 16p12.2-p11.2 deletion of approximately 7.7 Mbp^47^. The deletion carriers in these families presented with a diverse set of neurodevelopmental and psychiatric phenotypes (**Fig. 1B**), as has been previously described for this deletion^24,26^. We used hifiasm^48^ to generate haplotype-phased assemblies from all 19 individuals and further analyzed phased whole genome assemblies from an additional 293 individuals from HPRC^43^ and HGSVC^44^. Restricting our analysis to genomes that had the entire 16p12.2 region assembled in a single contig and at least 50 kbp from the ends of the contig, we identified 570 haplotypes that passed quality-control filters. These haplotypes represented diverse ancestries and included deletion haplotypes from seven individuals (**Fig. 1C, Table S1**).

We found significant diversity in the total length and sequence configuration of the 16p12.2 haplotype across individuals, beyond the S1 and S2 haplotypes originally described for this region^20^ (**Fig. 2A-B**). There was a 1.34-fold difference in the size of the 16p12.2 haplotype (ranging from 2.13 to 2.85 Mbp, excluding deletion haplotypes) across individuals, primarily resulting from variation in the SD blocks (**Fig. 2A, Table S1**). We further identified additional size variation in BP3 (range from 153 to 688 kbp) and, to a lesser extent, in BP2 (range from 200 to 547 kbp) beyond that previously described by others^27^. For example, East Asian (EAS) haplotypes showed 1.9-fold increase in size variability (standard deviation, 76 kbp) at BP3 than African (AFR) haplotypes (standard deviation, 41 kbp) (**Fig. 2B, Table S2**).

**Figure 2.**
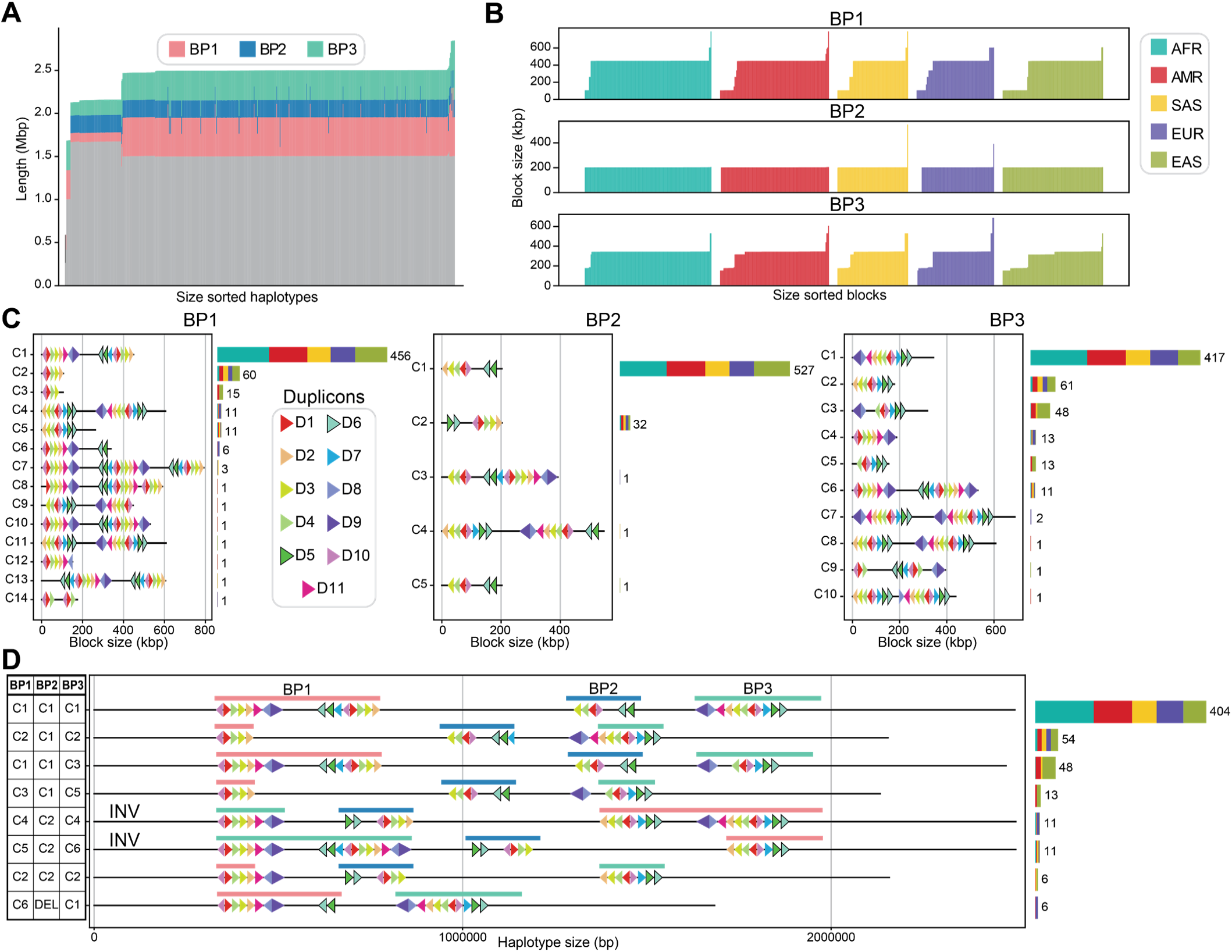
Haplotypes at 16p12.2 segmental duplication blocks. **(A)** Length of SD blocks (BP1, pink; BP2, blue; BP3, teal) and other sequence (grey) in the 16p12.2 locus in all haplotypes. **(B)** Total length of SD blocks identified in all haplotypes, colored by population. Size ranges, mean, median, and standard deviation are provided in Table S2. **(C)** Identified duplicon configurations at BP1 (left), BP2 (center), and BP3 (right). We note that one configuration passed initial QC, but the identity of the SD blocks could not be determined, so it has been excluded from these analyses (total analyzed n=569). Further, all 16p12.1 deletion haplotypes (n=7) do not have BP2, so n=562 haplotypes were analyzed for BP2 analyses. Additionally, the deletion haplotype of PC_326 also does not contain BP3, so n=568 haplotypes were analyzed for BP3 analyses. **(D)** Example 16p12.2 haplotypes showcasing the most common SD block architecture. Only haplotype configurations present in at least six haplotypes are shown. **(C-D)** Colored triangles show the order and orientation of duplicons. Bar plots show the number of haplotypes belonging to each cluster/configuration, colored by population.

Within these SD blocks, we identified duplicons of ∼20 kbp that occurred in multiple copies across the 16p12.2 locus in CHM13. We found 11 such duplicons that range in size from 18 to 35 kbp (**Fig. 2C**, **Table S3**). Notably, two duplicons, D5 (∼20 kbp) and D6 (∼21 kbp), are present at both BP1 and BP2 in CHM13 in a direct orientation and may mediate NAHR events contributing to the deletion. (**Fig. 2C**). To further define the organization of each SD block across samples, we calculated the Jaccard distance between all pairs of SD configurations at each block and performed hierarchical clustering on these distances to identify clusters of samples with similar architecture (**Fig. S1**). We identified 14 clusters at BP1, five clusters at BP2, and 10 clusters at BP3 (**Fig. 2C**). The most common cluster at each block (designated C1) represented the vast majority of SD architecture (80.1% of BP1, 93.8% of BP2, and 73.4% of BP3), although other clusters were observed at appreciable frequencies. Further, the SD clusters were not evenly distributed across ancestry groups. For example, the most common architecture at BP3, BP3-C1, was overrepresented in AFR and depleted in EAS haplotypes (Chi-squared test, p=5.87×10^-4^) and the second most common configuration at BP1, BP1-C2, was overrepresented in EAS and South Asian (SAS) haplotypes and depleted in AFR haplotypes (Chi-squared test, p=0.006) (**Table S4**). These clusters at each block combined to create 21 unique 16p12.2 haplotype configurations (**Fig. 2D**, **Fig. S2**, **Table S5-S6**). Interestingly, some haplotype configurations revealed inversions at the 16p12.2 locus (**Fig. 2D**). The most common clusters at BP1 and BP2, BP1-C1 and BP2-C1, have the D5 and D6 duplicons in the direct (NAHR-predisposing) orientation, and 79.8% of the 16p12.2 haplotypes (454/569) have this configuration (**Fig. 2D**, **Table S5**). Conversely, some of the less common haplotypes at BP1, including BP1-C2 and BP1-C3, do not contain the D5 and D6 duplicons and would thus be protective against NAHR and the 16p12.1 deletion (**Fig. 2C-D**). Only 13.7% of non-deletion haplotypes (77/562) have protective arrangements of D5 and D6 (**Fig. 2C-D**, **Table S5**).

### D5 and D6 duplicons mediate the 16p12.1 deletion

We next assessed how the SD architecture in the 16p12.2 region may predispose to the 16p12.1 deletion. The deletion haplotypes made up unique clusters at BP1 (BP1-C6 and BP1-C14) and lack BP2, suggesting that the deletion creates new SD architecture in the region (**Fig. 3A**). In fact, the structure of BP1-C6 indicates a recombination event between BP1-C1 and BP2-C1 occurring around D5 and D6 (**Fig. 3B**). Notably, the deletion haplotype in PC_326, the proband with the 16p12.2-p11.2 deletion, showed distinct SD architecture (BP1-C14) than that observed with the other deletion haplotypes (**Fig. 3A**), indicative of a different rearrangement in this proband. We also examined the 16p12.2 locus architecture in both alleles from the parent of origin of a proband with a *de novo* deletion (FNC_319) (**Fig. 3C**). This parent was homozygous for the most common configurations at each SD block (i.e., BP1-C1, BP2-C1, and BP3-C1), and this haplotype arrangement positions the D5 and D6 duplicons within BP1 and BP2 in a direct orientation, necessary to mediate NAHR. We further evaluated NAHR signatures in a proband with a *de novo* 16p12.1 deletion (PC_315) by assigning 16p12.2 locus reads from the proband to both haplotypes in the parent of origin and identifying recombination events where the assigned reads switched from one parental haplotype to the other (**Fig. 3D**). We found multiple recombination events, including an event localized around the D5 duplicon which likely represents the NAHR event that resulted in the deletion in this proband (**Fig. 3D**).

**Figure 3.**
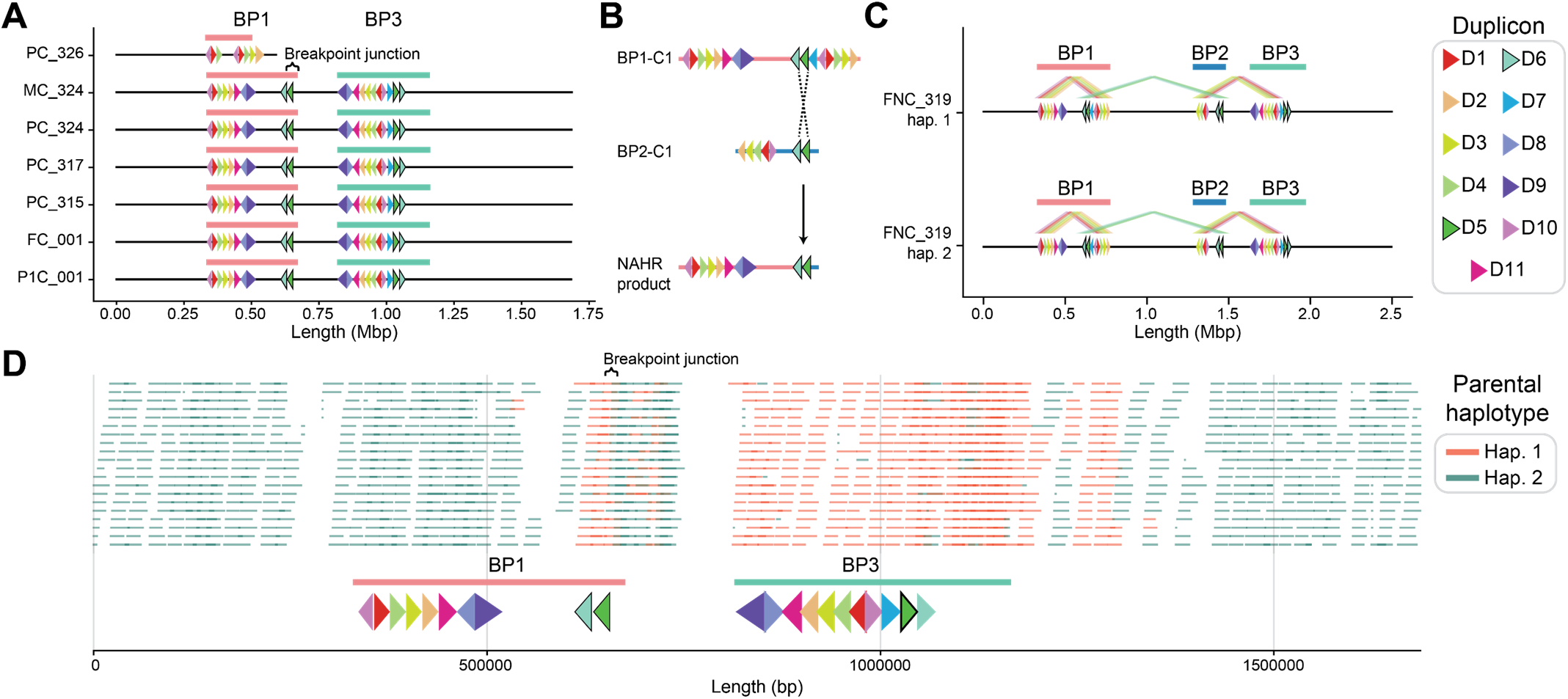
16p12.1 deletion haplotypes and non-allelic homologous recombination. Colored triangles indicate duplicon order and orientation for **(A)** seven 16p12.1 deletion haplotypes, **(B)** a schematic of NAHR between BP1-C1 and BP2-C1 at D5, **(C)** haplotypes in a parent of origin for a *de novo* 16p12.1 proband, and **(D)** deletion haplotype of a PC_315. In **(C)** lines above duplicons highlight pairs of duplicons in the direct (deletion-predisposing) orientation. **(D)** (Top) Reads from the 16p12.1 deletion haplotype in PC_315 assigned to haplotype 1 (orange) or haplotype 2 (green) in the parent of origin, FNC_315. Uninformative reads are not shown. (Bottom) Duplicon order and orientation for the 16p12.1 deletion haplotype in PC_315. In all panels, pink lines, BP1 boundaries; blue lines, BP2 boundaries; teal lines, BP3 boundaries.

### Deletion breakpoints are highly variable across families

Given strong evidence that the deletion is mediated by the D5 and D6 units in the SD blocks flanking the deletion region, we next sought to refine the breakpoints in samples with the deletion using PAV^44^ (**Fig. 4A, Table S7**). We found that the size of the deletion was much larger (approximately 815 kbp) than originally described (520 kbp)^12^ (**Fig. 4A**). This is likely due to both the improved accuracy of long reads for breakpoint detection and the improved quality of the T2T reference genome for highly repetitive regions^49^. Comparing breakpoints across the eight samples with 16p12.1 deletions showed that, while all breakpoints localized around D5 and D6, the deletion breakpoints varied by as much as 20 kbp across families (**Fig. 4A**). Furthermore, three families had partial deletions of *OTOA*, a gene associated with autosomal recessive hereditary hearing loss^50^. Notably, one proband with partial *OTOA* deletion, P1C_001, presented with hearing loss. In contrast to the variability across families, breakpoints in related samples were identical, suggesting that deleted alleles are stable across generations. Additionally, we were able to localize deletion breakpoints in the proband with the 16p12.2-p11.2 deletion to two copies of duplicon D10 in 16p12.2 and 16p11.2 (**Table S7-8**), suggesting that D10 mediated this deletion and resulted in the unique BP1 architecture in this proband (**Fig. 3A**).

**Figure 4.**
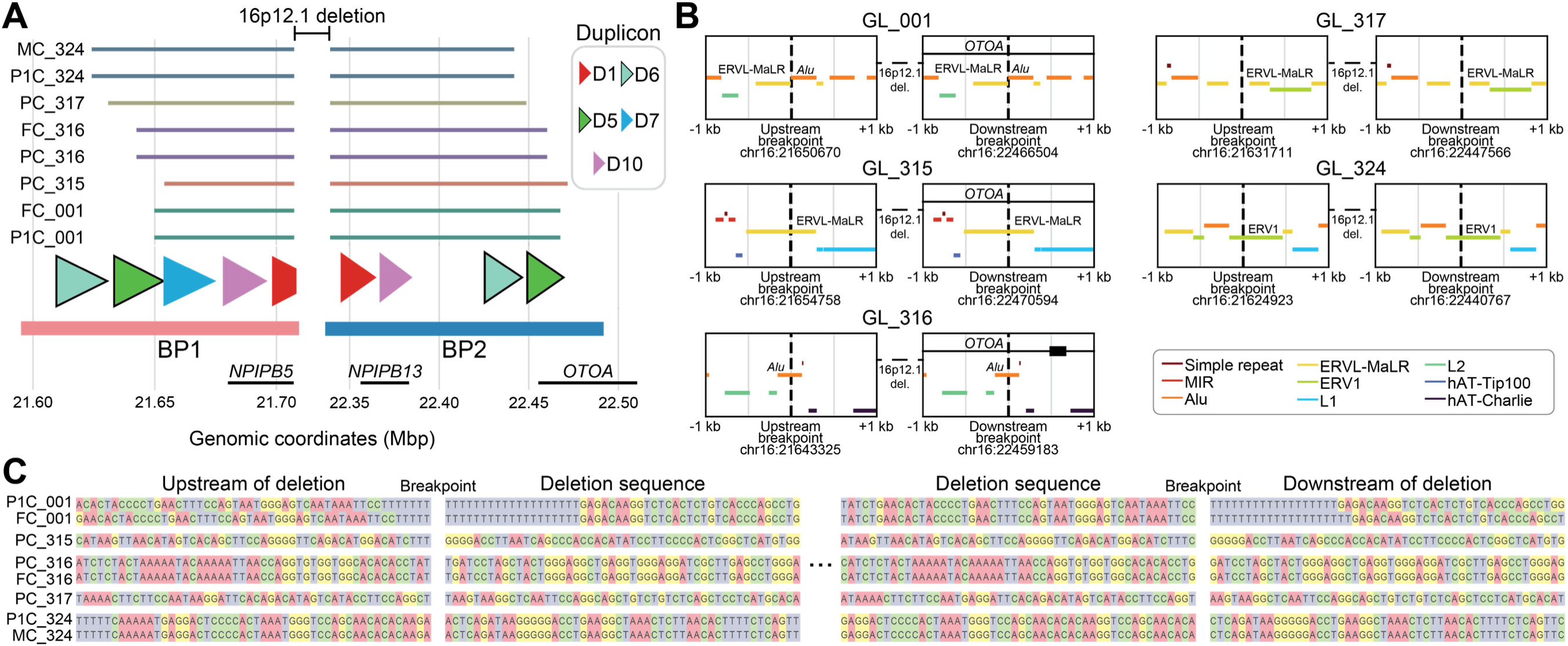
16p12.1 deletion breakpoints and local sequence. **(A)** 16p12.1 deletion breakpoints in eight individuals mapped to the CHM13-T2T reference genome. Colored triangles indicate duplicon order and orientation in CHM13-T2T, while pink bars indicate BP1 boundaries and blue bars indicate BP2 boundaries. Lines of the same color indicate related individuals. **(B)** 16p12.1 deletion breakpoints in five 16p12.1 deletion families and high copy repeat elements identified on CHM13-T2T within 1,000 bp of the breakpoints. Vertical dashed lines indicate deletion breakpoints, while horizontal colored lines indicate repeat elements. Horizontal black lines indicate intronic regions and black bars indicate exonic regions of *OTOA*. **(C)** (Left) 50 bp upstream of the deletion sequence and the first 50 bp of the deletion sequence in 16p12.1 deletion alleles. (Right) The last 50 bp of deletion sequence and the first 50 bp downstream of the deletion in 16p12.1 deletion alleles. In all panels, breakpoints of the atypical large deletion are not shown.

We then checked whether local sequence similarity at the variable breakpoints predisposes to the deletion across families. We first compared the high-copy repeat elements identified by Hoyt et al.^51^ in CHM13 against the breakpoints identified in each family (**Fig. 4B**). All families showed unique patterns of repeats around the deletion region, with breakpoints occurring within 100 bp of a repeat element (**Table S9**). For instance, the breakpoint junctions in GL_316 and GL_324 intersected an *Alu* element and an ERV1 element, respectively. Meanwhile, breakpoint junctions in GL_001 occurred between ERVL-MaLR and *Alu* elements. Further, although local sequence was highly similar at the upstream and downstream breakpoints in an individual, unique sequence signatures were observed in each family (**Fig. 4C**). These data suggest that while the D5 and D6 duplicons mediate the NAHR event in all families with the recurrent 16p12.1 deletion, the exact breakpoints and the flanking sequence architecture vary across families.

### Evolutionary toggling of the 16p12.2 locus

Given the structural diversity at 16p12.2 in humans, we next sought to investigate the evolutionary origin of this locus. We analyzed the CHM13 reference genome^42^ (HSA) and recent high-quality reference genomes of six ape species, including siamang (SSY), Bornean and Sumatran orangutan (PPY and PAB, respectively), gorilla (GGO), chimpanzee (PTR), and bonobo (PPA)^45^, and rhesus macaque (MMU)^46^. Comparison of sequence architecture across the primate species suggested extensive rearrangement of the 16p12.2 syntenic region (∼2.5 Mbp in humans) over evolutionary time, as reported previously for the p-arm of chromosome 16^45^ (**Fig. 5A-B, Table S10**). We first observed that ∼277 kbp of sequence containing multiple protein-coding genes at the distal end of the locus has translocated in orangutans to ∼1.5 Mbp upstream of the region (**Fig. 5B**). We further found the entire syntenic region to be inverted in SSY, PPY, and PAB relative to T2T-CHM13, but not in MMU. This suggests that the locus inverted once 19 to 25 Mya (million years ago) after the divergence of macaque and inverted a second time 11 to 19 Mya after the divergence of orangutan (**Fig. 5C**). Further, the unique regions between the SD blocks (interstitial region, I1 between BP1 and BP2, and I2 between BP2 and BP3) were in the same orientation in SSY, PPY, PAB, GGO, and human. However, both I1 and I2 were inverted in MMU, I1 was inverted in both PTR and PPA, and I2 was inverted in PPA, although previous work has shown that the I2 inversion is polymorphic in PPA^52^. This suggests that, separately from the whole 16p12.2 locus inversion, I1 and I2 inverted in the common ancestor after the divergence of orangutan, I1 inverted back in the PTR and PPA lineage, and I2 inverted back only in the PPA lineage (**Fig. 5C**). Notably, the occurrence of interstitial inversions at this locus coincides with the rapid burst of SDs in this region between the divergence of orangutan and gorilla^8^ (∼11 to 19 Mya) (**Fig. 5B**), suggesting that the emergence of these blocks may have facilitated inversions at this locus. These results are also reminiscent of other recurrent CNV loci, such as 17q21.31^53^ and 16p11.2^52^, which have been shown to “toggle” between orientations over evolutionary time.

**Figure 5.**
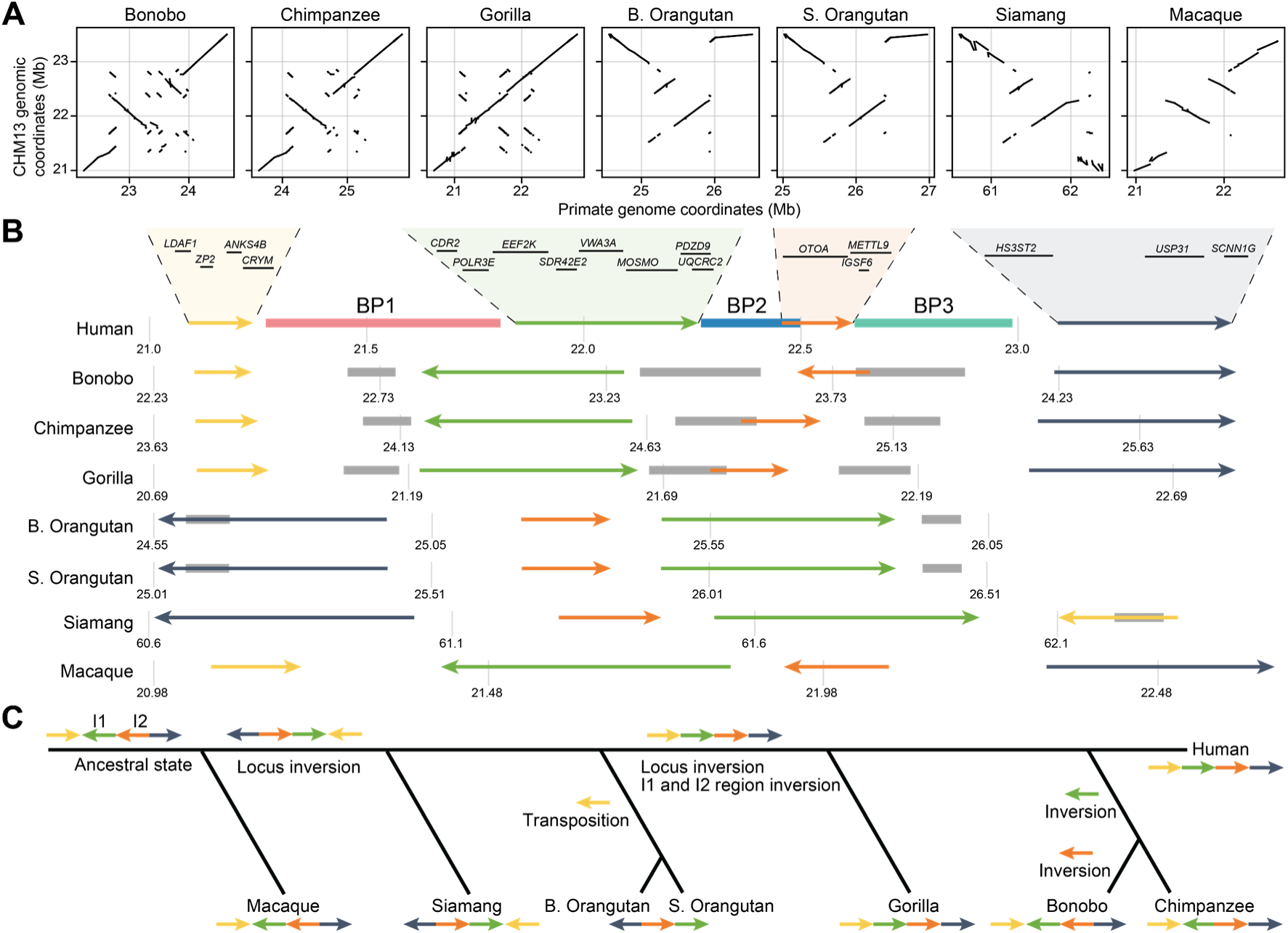
Evolutionary history of the 16p12.2 locus. **(A)** Dot plots of the 16p12.2 syntenic region in seven primate species compared to CHM13-T2T. **(B)** Order and orientation of genic regions in the 16p12.2 region in primate species. Colored arrows indicate genic regions. Colored bars indicate BP1-3 in CHM13-T2T, while grey bars indicate SD blocks in other species. **(C)** Schematic of the most parsimonious evolutionary history of inversions within the 16p12.2 locus in primates.

Given the history of inversions at the 16p12.2 locus, we next sought to examine human haplotypes with inversions, some of which have been reported previously^21,52,54^. Using PAV, we identified inversions in 32 of the 570 16p12.2 haplotypes ranging in size from ∼472 kbp to ∼1.44 Mbp (**Fig. 6A-B, Table S11**). While breakpoints were variable across samples, most inversions (n=26) were between BP1 and BP3, five were between BP1 and BP2, and a single inversion was found between BP2 and BP3. One sample (NA20503) was homozygous for the BP1-BP3 inversion, and another sample (HG00232) was compound heterozygous for inversion of both BP2-BP3 and BP1-BP3.

**Figure 6.**
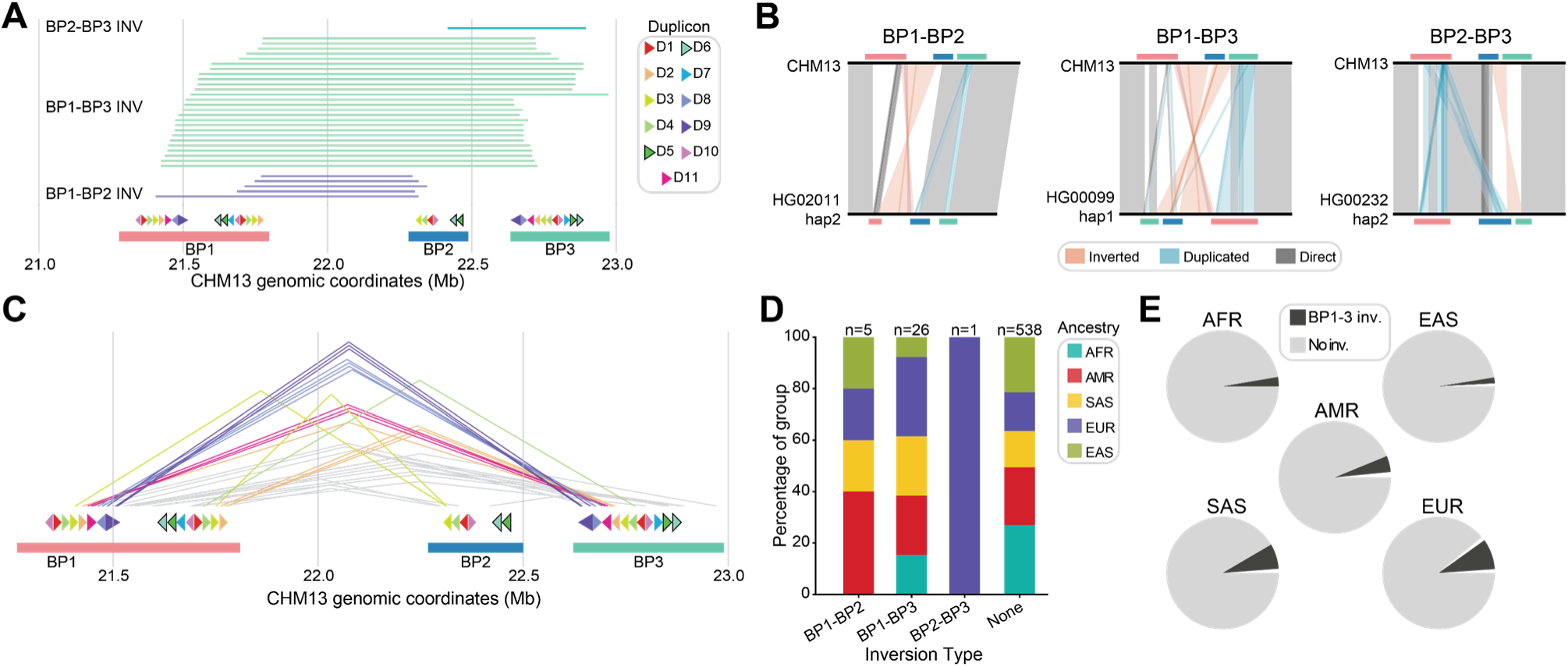
16p12.2 inversion in humans. **(A)** Inversion breakpoints identified in human haplotypes relative to CHM13. Colored triangles indicate duplicon order and orientation in CHM13-T2T, while pink bars indicate BP1 boundaries, blue bars indicate BP2 boundaries, and teal bars indicate BP3 boundaries. Colored lines represent inversion regions with breakpoints in the same SD blocks. **(B)** Examples of inversion haplotypes mapped to CHM13. Orange lines, inverted regions; blue lines, duplicated regions; grey lines, single-copy and directly oriented regions. **(C)** Inversions colored by the duplicons their breakpoints intersect. Grey lines indicate inversions that overlap no duplicons or where each breakpoint maps to a distinct duplicon. **(D)** Percentages of haplotypes with and without inversions colored by ancestry group. **(E)** Proportion of haplotypes with BP1-BP3 inversions within each ancestry group.

While the 16p12.1 deletion is mediated by the D5 and D6 duplicons, duplicons mediating inversions were more variable (**Fig. 6C**, **Table S11**). We found several instances in our samples where both breakpoints mapped to the same duplicon across blocks: BP1-BP3 breakpoints mapped to D2 (n=4), D4 (n=1), D8 (n=4), D9 (n=3), and D11 (n=3), while BP1-BP2 breakpoints mapped to D3 (n=2). Although small sample sizes limit statistical power, BP1-BP3 inversions were not evenly distributed across ancestries (Chi-squared test, p=0.09) (**Fig. 6B-C**). For instance, European (EUR) haplotypes were 31% (8/26) and SAS haplotypes were 23% (6/26) of the observed BP1-BP3 inversions, despite representing only 16% and 15% of the cohort, respectively. We found the prevalence of BP1-BP3 inversions to be 2.7% in AFR, 4.6% in Admixed American (AMR), 1.7% in EAS, 8.8% in EUR and 7.2% in SAS haplotypes (**Fig. 6C**), while the other inversions occurred at much lower frequencies. While the inversions present in other primate species were also found in human 16p12.2 haplotypes, such as BP1-BP2 (I1) inversion in PTR and PPA and BP2-BP3 (I2) inversion in PPA (**Fig. 5**), the most common inversion identified in humans, BP1-BP3 inversion, was not observed in the other primates.

### Incomplete lineage sorting and lineage-specific expansions in 16p12.2 SDs

We next sought to investigate the history of the SD blocks at this locus. The amount of unique sequence has remained relatively constant, apart from the translocation in orangutan (**Fig. 7A**, **Fig. 5**). However, the amount of duplicated sequence has increased dramatically since the divergence of orangutan, with >350 kbp of additional duplicated sequence in GGO (∼526 kbp) compared to SSY (∼163 kbp) (**Fig. 7A**), in agreement with past analysis of SD expansions in primates^8^. PTR (∼585 kbp), PPA (∼710 kbp), and CHM13-T2T (∼879 kbp) all show increased amounts of duplicated sequence compared to GGO (**Fig. 7A**). Because the amount of duplicated sequence was variable in humans (**Fig. 2A-B**), we compared all human haplotypes to the other primates and found that the average amount of duplicated sequence in humans (excluding deletion haplotypes) was ∼848 kbp. However, 74 haplotypes showed ≤646 kbp of duplicated sequence, ∼64 kbp less than the bonobo genome.

**Figure 7.**
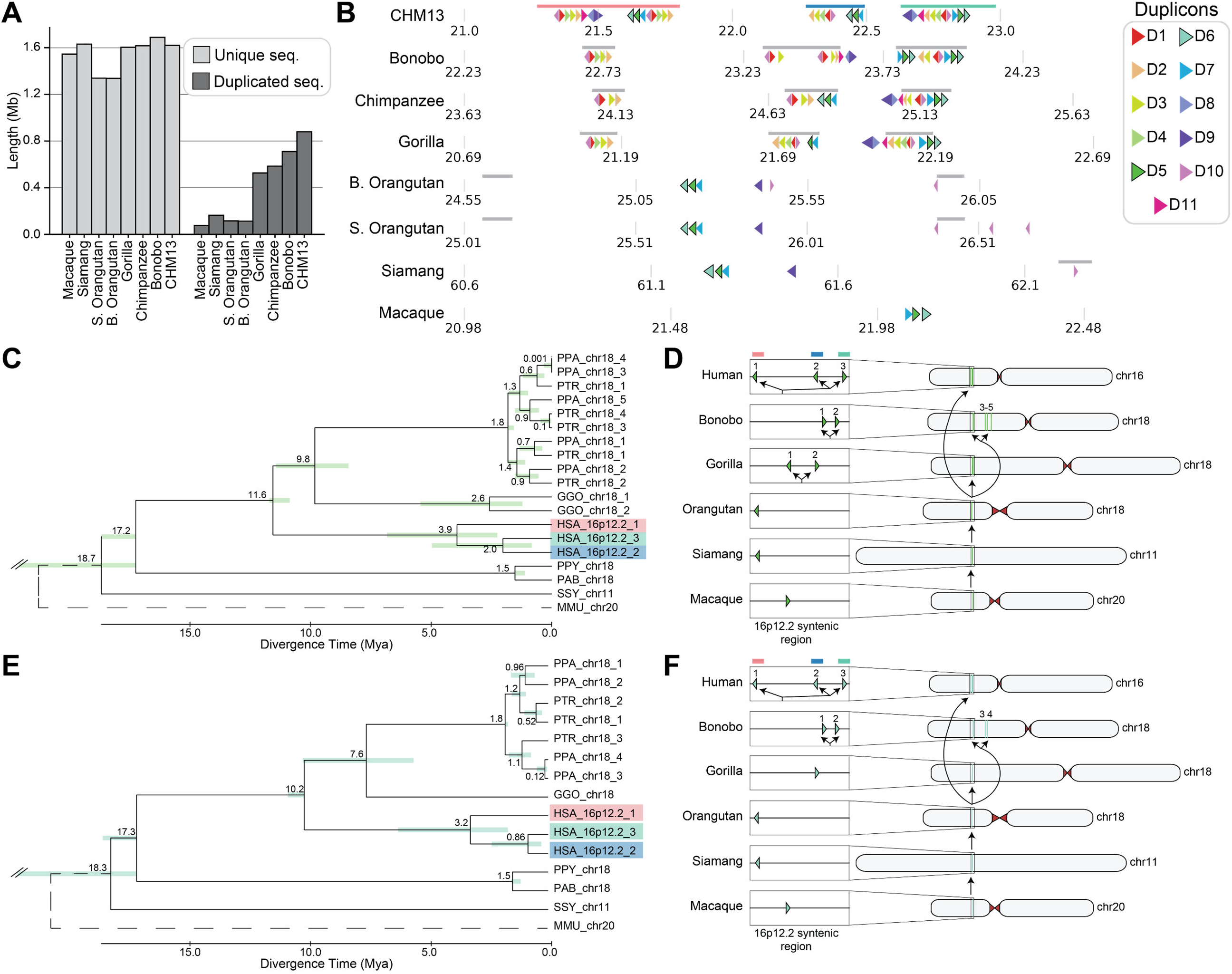
Evolution of segmental duplication blocks. **(A)** Length of unique (light grey) and duplicated (dark grey) sequence in primate genomes. **(B)** Colored triangles indicate duplicon order and orientation in primate genomes. Colored bars indicate BP1-3 in CHM13-T2T, while grey bars indicate SD blocks in other primate species. **(C, E)** Phylogenetic tree of duplicon D5 (**C**) and D6 (**E**) duplications in primate species with macaque outgroup. Shaded bars indicate the 95% confidence interval (CI) for the divergence time. The upper bound of the CI for the divergence of siamang is clipped but extends to 31.8 Mya for D5 and 36.0 Mya for D6. **(D, F)** Schematic detailing the evolutionary history of the D5 (**D**) and D6 (**F**) duplicons. In **D** and **F**, numbers represent the copies shown in **C** and **E**, such that “1” in the human haplotype represents HSA_16p12.2_1. In **(C-F)**, colored boxes around the human paralogs correspond to the SD blocks they lie within (BP1, pink; BP2, blue; BP3, teal).

Given the variability in duplicated sequence, we assessed how the 11 duplicons we identified in human haplotypes were conserved across primates. We mapped duplicons identified in CHM13 to the whole genome of each primate species (**Fig. 7B**, **Table S12**) and found that the patterns of mapped duplicons were indicative of extensive duplication and rearrangement in the region. We were able to identify duplicons which flank inverted loci in opposite orientations in non-human primates, which may have mediated the inversions observed between species. D1, D2, D3, and D10 in PPA and PTR flank I2 and D1, D2, D3, D4, and D10 in GGO flank I1 (**Fig. 7B**). Additionally, many of the duplicons showed ancient origins in the primate lineage, with six duplicons (D2-D7) present in rhesus macaque and most (D2-D11) present in siamang (**Fig. 7B**, **Table S12-13**). Duplicons varied significantly in copy number across species and mirrored the increases in duplicated sequence across species, with higher copy numbers in the African great apes compared to other species (**Table S13**). One duplicon, D9, had only duplicated in the human lineage, with a single copy in all other apes (**Table S12-13**). This proliferation was not restricted to the 16p12.2 locus; many duplicons had paralogs at other loci or chromosomes. For example, D1, D2, D3, D4, D8, D10, and D11 all had paralogs in humans at 16p11.2. Duplicon D10 was particularly prolific, with more than 20 paralogs in all great apes and paralogs on multiple chromosomes in those species (**Table S12-13**).

We further examined the evolutionary history of these duplicons by generating phylogenetic trees for each duplicon (**Fig. 7C-F**, **Figure S2-10**). These phylogenies showed evidence of incomplete lineage sorting (ILS). For example, the phylogenies of D5 and D6 showed GGO and PPA and PTR orthologs cluster more closely than HSA orthologs (**Fig. 7C**, **Fig. 7E**). This discrepancy between the species phylogeny and the observed duplicon phylogeny is indicative of ILS, a phenomenon that is common in primate genomes^55^ and has been observed for other SDs^56^. In fact, most (8/11) duplicons had phylogenies consistent with ILS, which was primarily seen in the common ancestor of the African great apes, with a clear separation of siamang and orangutan lineages for most duplicons (**Fig. 7C-F**, **Fig. S2-10**).

The phylogenetic trees of D5 and D6 also showed evidence of lineage-specific expansions (LSE) (**Fig. 7C**, **Fig. 7E**). For example, D5 was present only once in MMU, SSY, PPY, and PAB (**Fig. 7C**). However, it was present at two copies in GGO, three in HSA, and five in PPA and PTR and phylogenetic tree reconstruction appeared to show that D5 duplicated independently in each lineage (**Fig. 7C-D**). In the human lineage, D5 appeared to have duplicated once to create the BP1 copy approximately 3.9 Mya and duplicated a second time approximately 2.0 Mya to create the BP2 and BP3 copies (**Fig. 7C-D**). From the human haplotypes, we know these duplications are polymorphic in the lineage; D5 was present in only 86.5% (492/569) of BP1 haplotypes and 97.9% (556/568) of BP3 haplotypes but was present in 100% (562/562) of BP2 haplotypes (**Fig. 2C, Table S14**). This may suggest that the BP2 copy of D5 was the ancestral paralog of the duplicon shared with other primate species. A similar pattern of LSE was observed for D6, although it was present in fewer copies in GGO (one copy), PTR (three copies), and PPA (four copies) than D5 (**Fig. 7E-F**). The absence of D5 and D6 paralogs in a direct orientation on both sides of I1 in non-human primates suggests that only humans would be susceptible to the recurrent 16p12.1 deletion mediated by these duplicons (**Fig. 7B**). However, D2, D3, and D10 were directly oriented around the same region in PTR and PPA and could potentially predispose to similar rearrangements (**Fig. 7B**).

In phylogenies of all duplicons, including D5 and D6, we observed patterns of closer clustering within species rather than across species. In particular, the 16p12.2 locus paralogs are clustered together for all duplicons (**Fig. 7C-F**, **Fig. S2-10**). However, some duplications did appear to show ancestral origin, such as the copies of D2 and D3 on 16p12.3, which clustered more closely with orthologs in other African great apes than they did with paralogs in humans (**Fig. S3-4**). Across phylogenies where duplicon copies appeared at more than one locus, there was a consistent pattern where duplicon copies that are near one another clustered more closely than those that are farther away. Although these patterns may be evidence of LSE, they may also be evidence of interlocus gene conversion (IGC), which has been shown to occur extensively in SDs^57^. This presented two potential scenarios (**Fig. 8**). The first is that some of these duplicon copies represent ancient duplications that separated over time^58^ and the clustered paralogs represent more recent duplications that have yet to separate. The second is that many of these paralogs represent ancient duplications, but copies that remained close together “homogenized” via IGC^59^, leading to the observed clustering of paralogs. Indeed, the shared relative positioning of duplicons in the 16p12.2 locus across African great apes (**Fig. 7B**) supports the IGC hypothesis. While the phylogenetic trees of some duplicons clearly show LSE, such as the human-specific duplications of D5 and D6 into BP1, independent duplication events leading to the same patterns of duplicon position and orientation are unlikely, such as the copies of D5 and D6 in BP2 and BP3 in human and the other African great apes. Thus, we hypothesize that both LSE and IGC have occurred extensively in the 16p12.2 SD blocks.

**Figure 8.**
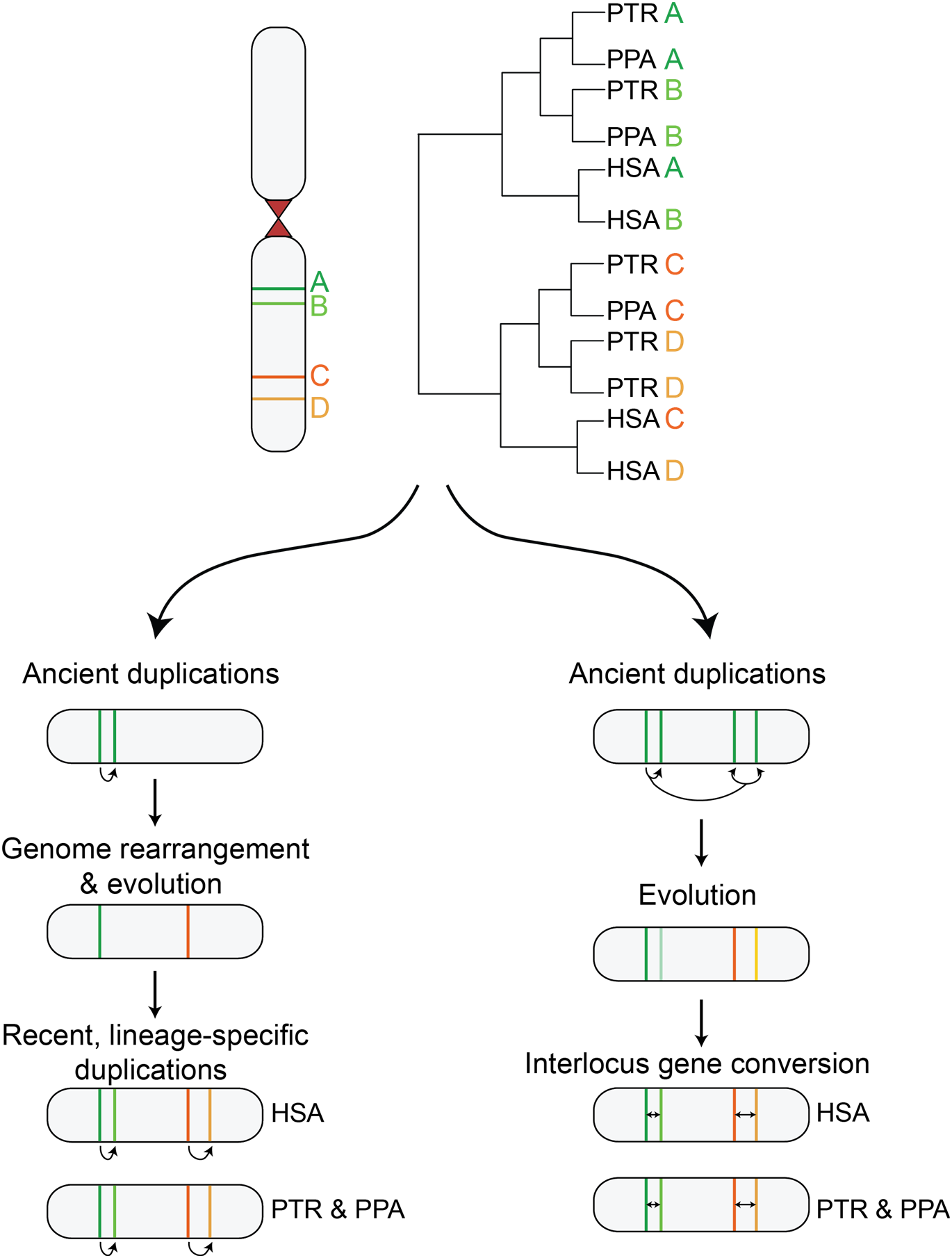
Models of lineage specific expansions and interlocus gene conversion in duplicon evolution. Two models for the evolutionary histories of duplicons within the 16p12.2 locus based on patterns of lineage-specific clustering. (Left) Paralogs at different loci are the result of ancient duplication events in the common ancestor of multiple ape species. Over time, these paralogs drifted apart and, independently in each lineage, duplicated again. (Right) Duplications at two distinct loci and the same locus are the result of ancient duplication events in the common ancestor of multiple ape species. Over time, these paralogs evolved separately, but those in the same locus underwent interlocus gene conversion in each lineage.

## DISCUSSION

We used high quality phased genome assemblies from diverse ancestries and telomere-to-telomere primate reference genomes to provide a comprehensive view of the 16p12.2 locus. This in-depth exploration of the locus revealed some key insights. We firstly identified more size diversity at the SD blocks than has been previously described^20,27^, identifying multiple haplotype configurations at each block (**Fig. 2**). These repeat regions exhibit extensive variability in size and structure, confirming and expanding upon previous work^27^. We further describe the variability in these configurations across diverse ancestries using the largest collection of samples to date. As reported previously^20,27^, EAS samples are enriched for BP1-C2, which is likely protective against the deletion. We also found an enrichment in BP1-C2 in SAS samples. These differences in haplotype structure have important implications for deletion risk, as populations depleted for BP1-C1 would have lower risk of NAHR. While we were able to examine structural differences across ancestries, we were not able to explore haplotype variability within ancestral groups. Additional investigations are required to determine if some populations within these groups are further enriched or depleted for protective haplotype structures.

We further explored the detailed architecture of these 16p12.2 haplotypes, describing 11 duplicons in the region. Past work proposed that the deletion was mediated by an ∼68 kbp repeat unit^27^ in BP1 and BP2. We refined this region to two ∼20 kbp repeat units, D5 and D6, and observed that, while deletion breakpoints localized around these duplicons in all families, they vary across families in location, local sequence, and flanking landscape of high copy repeats (**Fig. 4**). This suggested that while D5 and D6 likely mediate the deletion, NAHR events can occur anywhere within these duplicons. Further investigation of more families will determine if any breakpoints are recurrent across multiple families, or if all families show unique breakpoints. Recent studies have suggested that this variability is not unique to the 16p12.1 deletion. Other CNV disorders, including 22q11.2^36,60^ and 3q29 CNVs^37^, showed similar variability in breakpoints. Further, many CNV disorders, including the 16p12.1^22^ and 22q11.2^61^ deletions, are characterized by phenotypic variability^62^. While deletion breakpoint variation is unlikely to be the primary cause of the variable phenotypes associated with these CNVs, it may account for some of the observed phenotypic heterogeneity across affected families. This may be particularly relevant in cases where CNVs affect additional genes, such as the families with partial *OTOA* deletions in our cohort (**Fig. 4A-B**). Notably, one of the three probands with partial *OTOA* deletion had impaired hearing and in a larger cohort of probands with 16p12.1 deletion^26^, 11.7% (9/77) presented with hearing loss. With increased samples sizes, we may be able to further characterize the full spectrum of breakpoint variability and describe the amount of phenotypic heterogeneity, particularly the incomplete penetrance of hearing loss phenotypes in 16p12.1 deletion carriers, attributable to variable breakpoints and partial *OTOA* deletions.

We also examined the evolutionary history of inversions of this locus. We observed that the region has undergone multiple inversions and rearrangements in the primate lineage (**Fig. 5**). It was previously suggested that the I2 interval was inverted in all African great apes compared to orangutan and macaque^20^, but by leveraging new telomere to telomere primate reference genomes we have shown that this region is in the same orientation in the reference genomes of orangutan, gorilla, chimpanzee, and human and inverted in bonobo and macaque (**Fig. 5**). We additionally found 32 cases of inversions in the human haplotypes, primarily between BP1-BP3 (**Fig. 6**), some of which have been reported previously^21^. These data suggest that the region is relatively unstable, toggling between orientations over evolutionary time, a rare feature in primate genomes. Porubsky et al. identified only 23 loci that exhibit this “toggling”, including 16p12.2 and 16p11.2^52^, likely mediated by the SDs that flank these regions. Accordingly, we found that I1 and I2 inversions were observed only in the African great apes, species with SDs flanking those regions. In humans, inversion breakpoints suggest that at least six duplicons within the SDs can mediate 16p12.2 locus inversions, compared to only two duplicons (D5 and D6) that mediate the 16p12.1 deletion.

We further examined the history of duplicons within the 16p12.2 SD blocks. Phylogenetic trees for each duplicon revealed ILS in the common ancestor of gorilla and human and numerous examples of LSE (**Fig. 7, Fig. S2-11**). In fact, the deletion-mediating D5 and D6 duplicons duplicated into BP1 only in the human lineage and this architecture is polymorphic in humans, contributing to differential risk for deletion across haplotypes. Evolutionary histories of all duplicons showed lineage-specific clustering of paralogs. While this pattern is consistent with LSE, it is also consistent with IGC between paralogs (**Fig. 8**). SDs are prone to IGC^63^, including those at other disease-associated loci, such as 22q11.2^64^. This process can obfuscate the history of duplication of these sequences and could be the cause of the lineage-specific clustering of duplicon copies we observed. Additional investigation will be needed to fully disentangle the history and effects of LSE and IGC in the 16p12.2 SD blocks.

While we have performed the most comprehensive characterization of the 16p12.2 locus to date, our study has limitations. Firstly, our analyses are reliant on accurate assemblies. We filtered for high-quality assemblies of the 16p12.2 region but even with long read technologies, highly repetitive sequences are challenging to assemble^65^ and inaccurate assemblies could confound our analyses. Additionally, our analyses of human haplotypes suggest that the 16p12.2 locus is highly variable, but our investigations in other primates are based on single reference genomes. While these are the most complete genomes available, additional variation at the 16p12.2 locus likely exists within primate populations. Indeed, a recent study showed that the orientation of the I2 region is polymorphic in bonobos^52^ and others have shown copy number variation across apes^66^, suggesting that additional polymorphism in the region is likely in other primates. Further studies using more genomes from non-human primates can determine if the variability at this locus is unique to humans or is also seen in other primates, particularly in other African great apes. Our current investigations into the evolutionary history of this locus showed the relationship between SD expansions in African great apes, the resulting genomic instability, and the haplotype diversity and disease risk in modern humans. Detailed study of the haplotype structure and evolutionary history conferring risk or protection against 16p12.2 rearrangements will provide more insights in understanding its risk for a range of neurodevelopmental outcomes.

## Data and code availability

All bioinformatic pipelines relating to data processing and analyses are available on GitHub (https://github.com/csmolen6136/16p12.2-Locus-Characterization).

## Acknowledgements

This work was supported by NIH R01-GM121907 and resources from the Huck Institutes of the Life Sciences to S.G. We are grateful to all of the families who provided samples and phenotypic information for this study, without whom this work would not have been possible. We would like to acknowledge the Human Pangenome Reference Consortium (BioProject ID: PRJNA730823) and the Human Genome Structural Variation Consortium and their funder, the National Human Genome Research Institute (NHGRI). We thank Karen Runkle (Penn State) and Serena Noss (Penn State) for their assistance recruiting families for this study and processing patient samples. We also thank Christian Huber, Heather Hines, and Dave Towes for their insights and helpful discussions on the analyses.

## Author Contributions

C.S. and S.G. conceived the idea and designed the experiments. C.S. performed all analyses. C.S. and S.G. wrote the manuscript.

## Declaration of Interests

The authors declare no competing interests.

## SUPPLEMENTARY FIGURES

**Figure S1.**
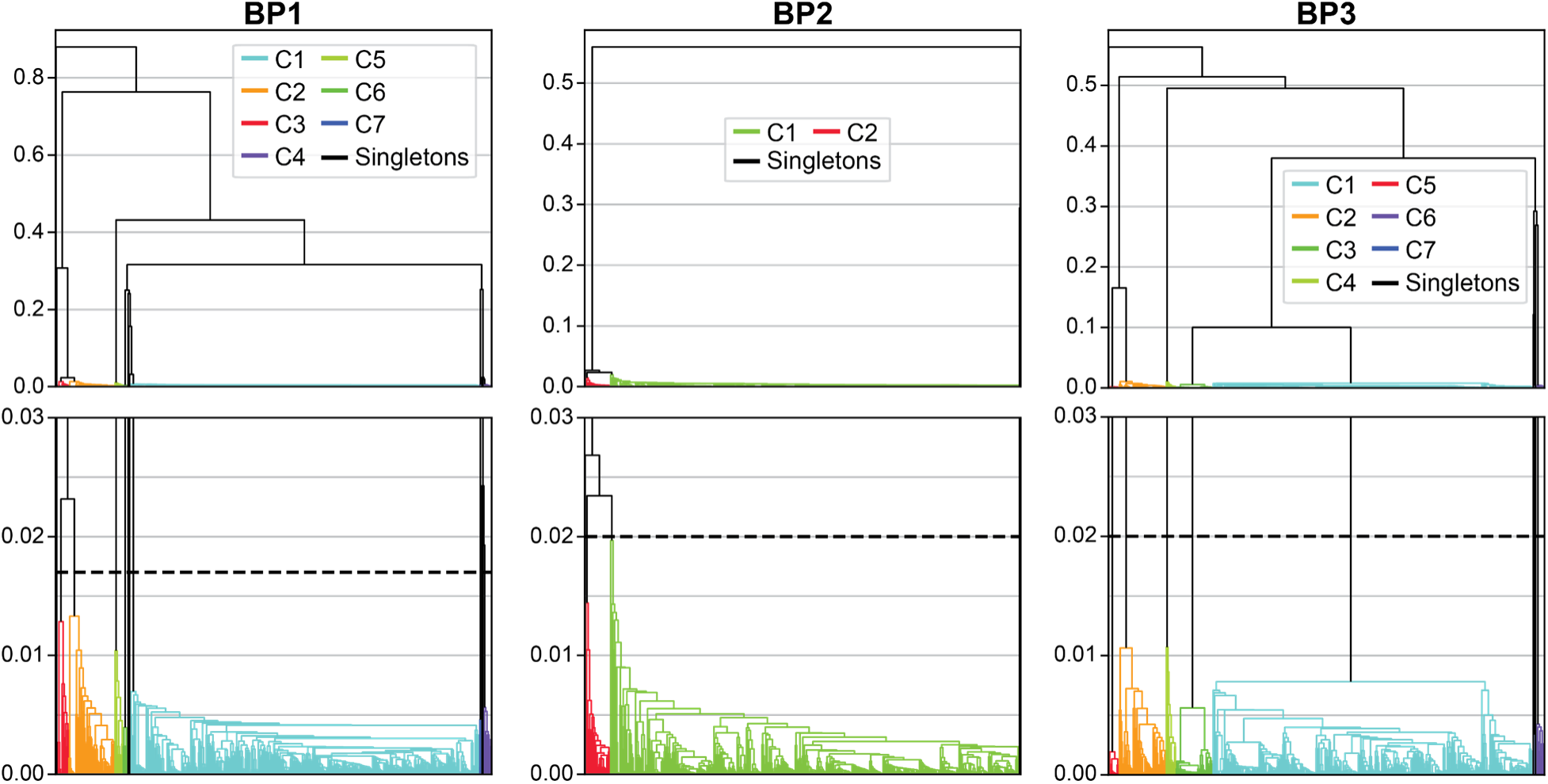
Hierarchical clustering of identified SD blocks. Hierarchical clustering of SD blocks in human haplotypes using Jaccard distance scores for BP1 (left), BP2 (center), and BP3 (right). Colored lines indicate samples belonging to the same block cluster. All singleton clusters are shown with black lines. Bottom panels show the same data as the top panels, zoomed in to highlight detail. Dashed horizontal lines indicate threshold used to determine group identity.

**Figure S2.**
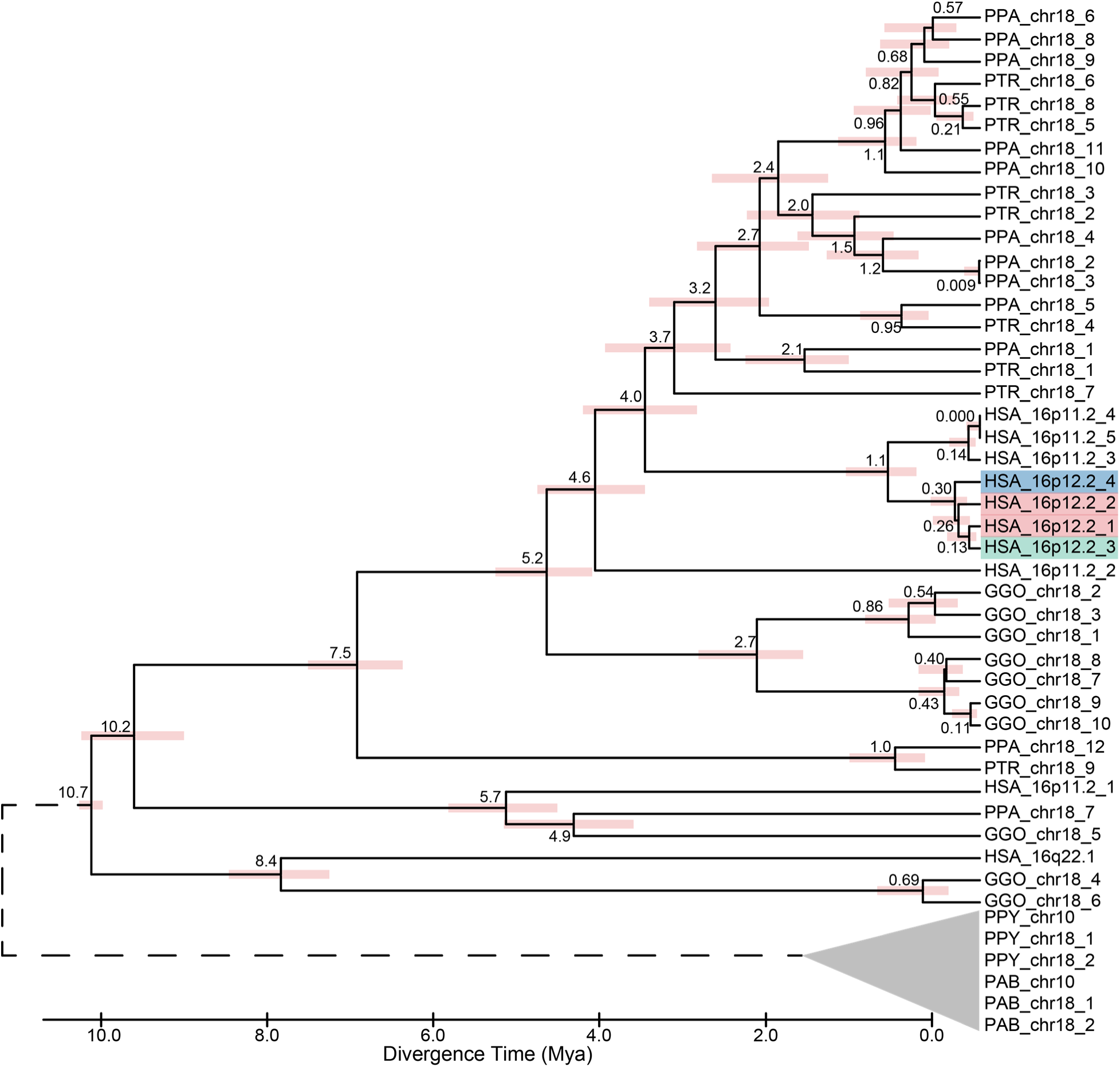
Phylogenetic tree for duplicon D1. Phylogenetic tree for D1. Dotted line indicates outgroup and colored bars indicate 95% confidence interval. Human 16p12.2 copies are shaded by the SD blocks they reside in: pink, BP1; blue, BP2; teal, BP3.

**Figure S3.**
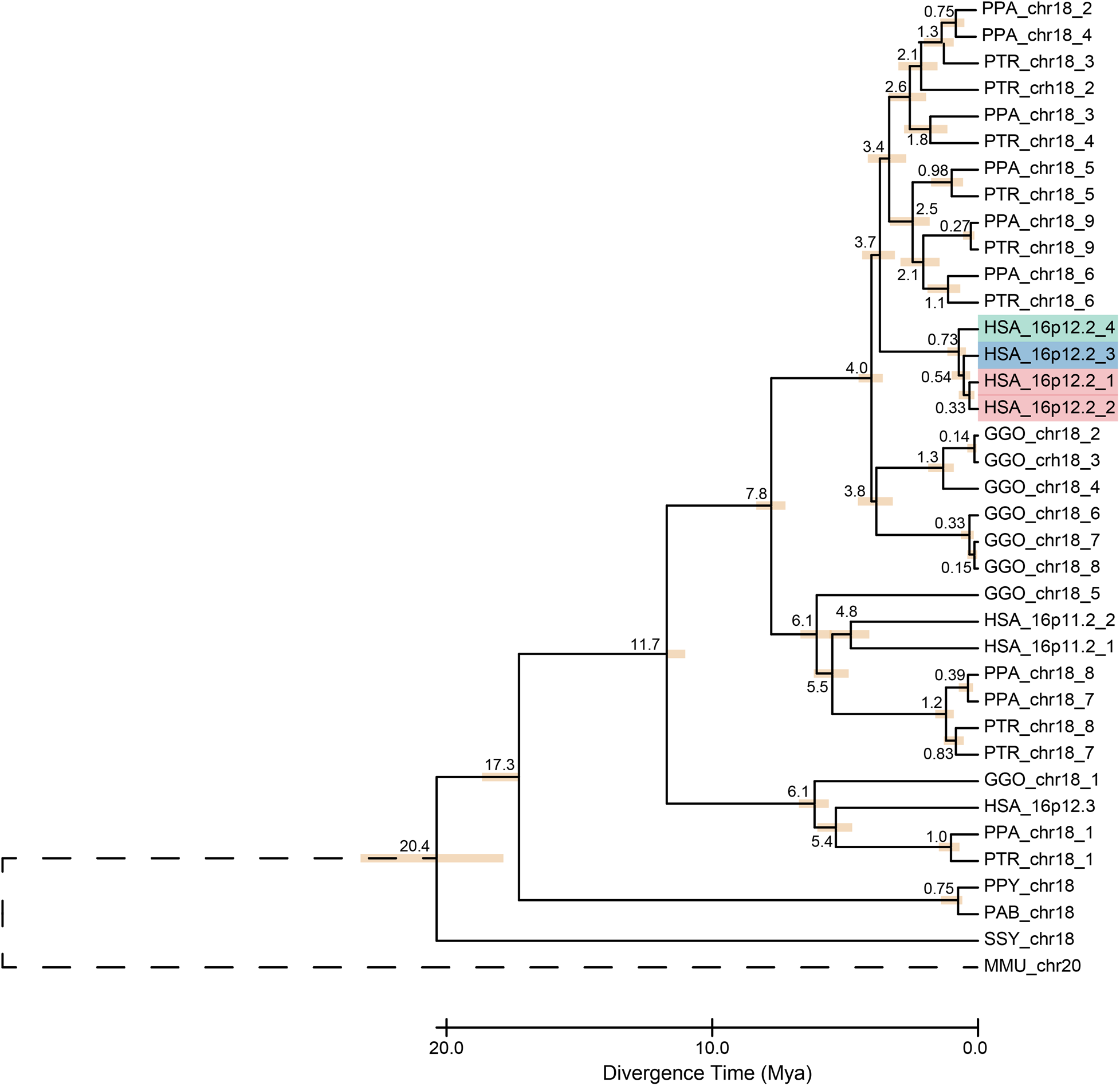
Phylogenetic tree for duplicon D2. Phylogenetic tree for D2. Dotted line indicates outgroup and colored bars indicate 95% confidence interval. Human 16p12.2 copies are shaded by the SD blocks they reside in: pink, BP1; blue, BP2; teal, BP3.

**Figure S4.**
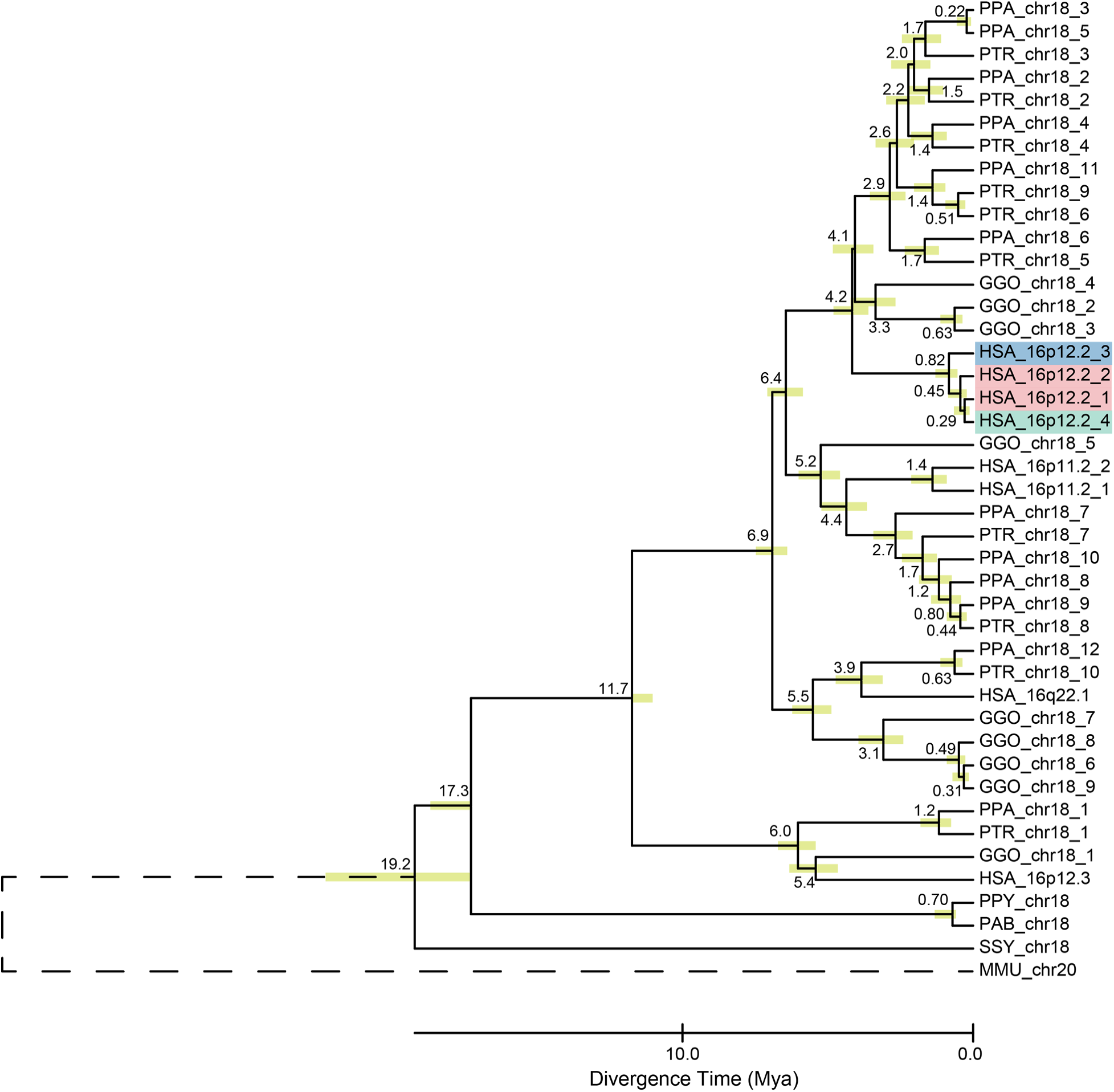
Phylogenetic tree for duplicon D3. Phylogenetic tree for D3. Dotted line indicates outgroup and colored bars indicate 95% confidence interval. Human 16p12.2 copies are shaded by the SD blocks they reside in: pink, BP1; blue, BP2; teal, BP3.

**Figure S5.**
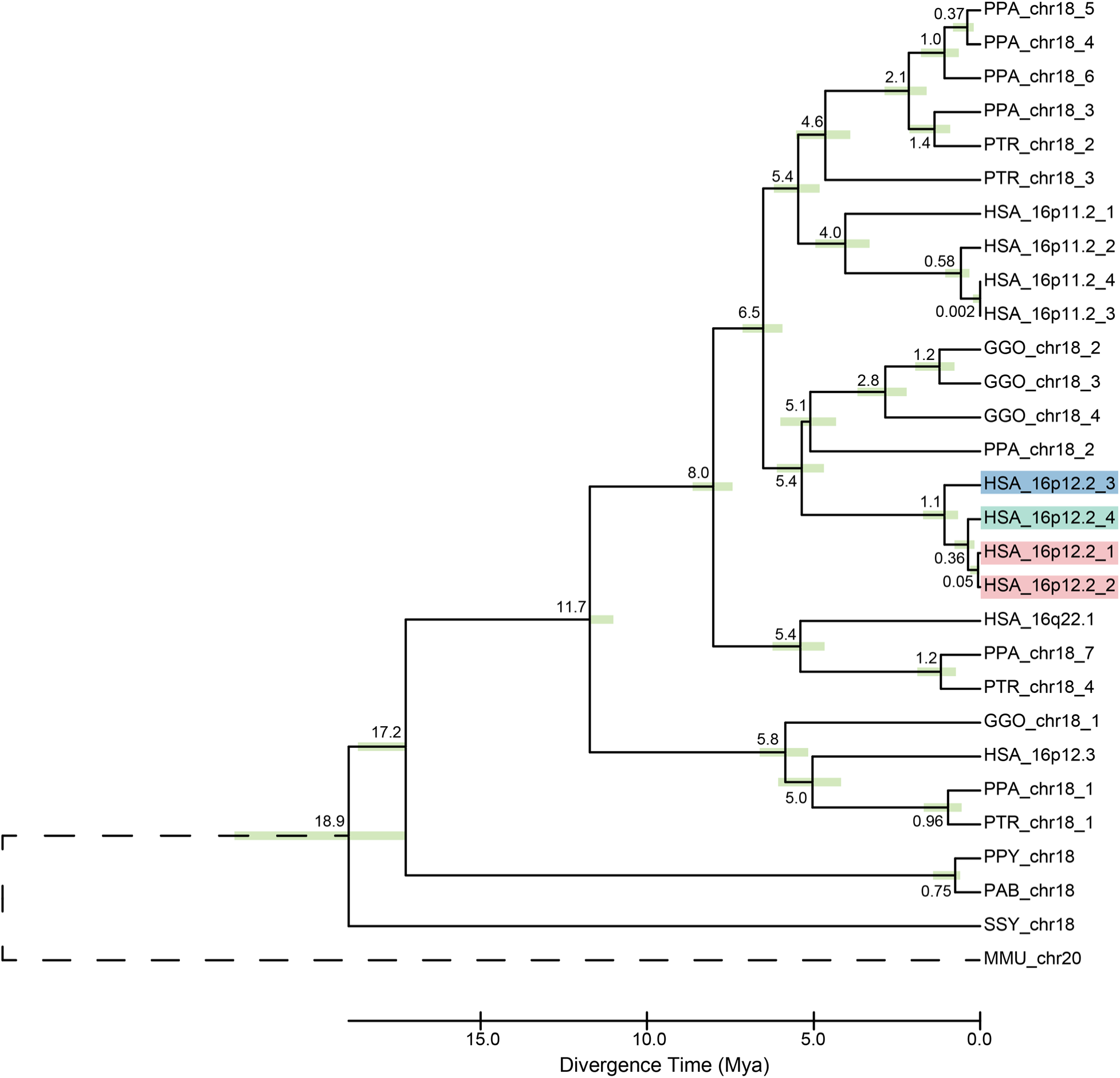
Phylogenetic tree for duplicon D4. Phylogenetic tree for D4. Dotted line indicates outgroup and colored bars indicate 95% confidence interval. Human 16p12.2 copies are shaded by the SD blocks they reside in: pink, BP1; blue, BP2; teal, BP3.

**Figure S6.**
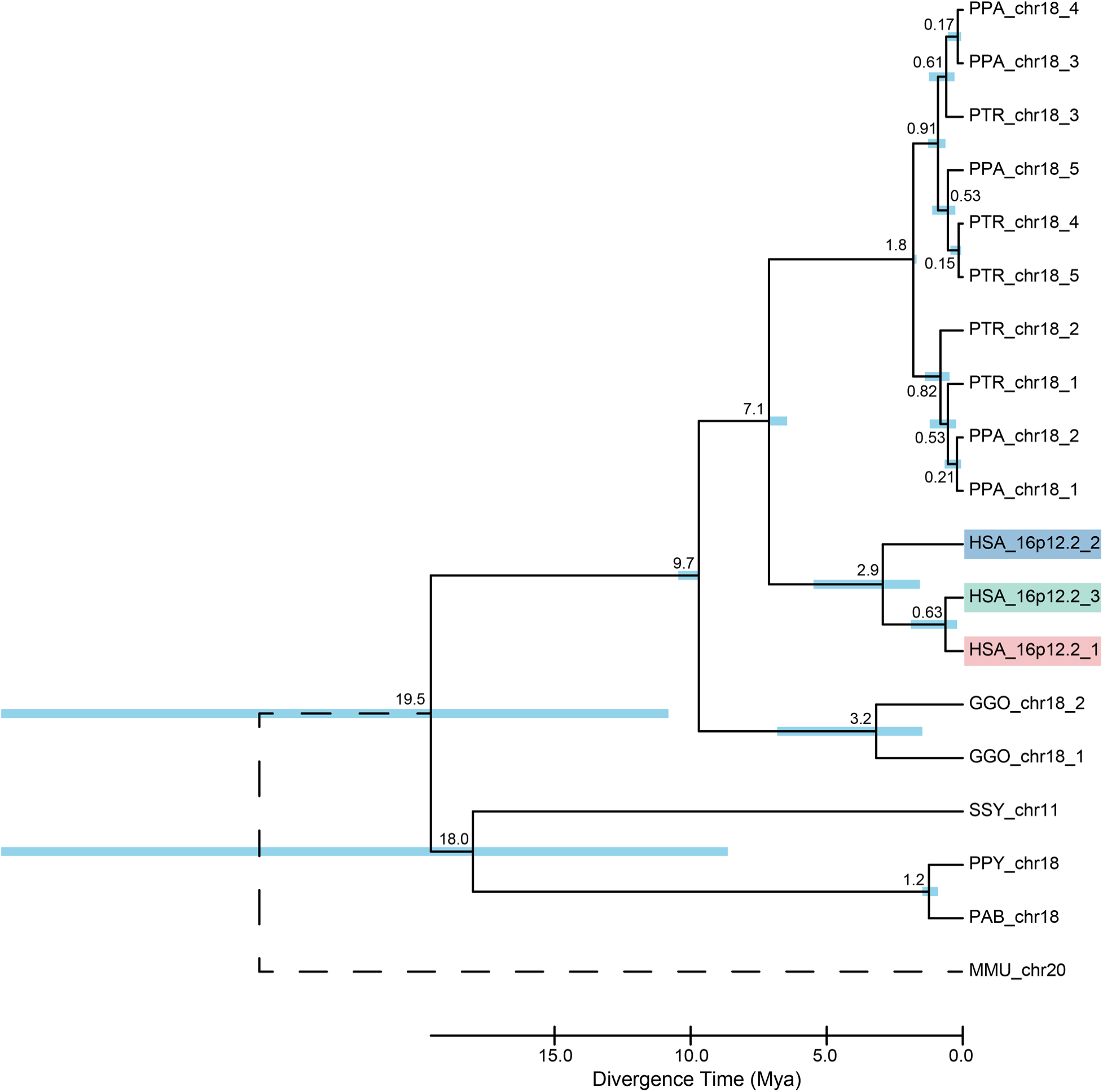
Phylogenetic tree for duplicon D7. Phylogenetic tree for D7. Dotted line indicates outgroup and colored bars indicate 95% confidence interval. Human 16p12.2 copies are shaded by the SD blocks they reside in: pink, BP1; blue, BP2; teal, BP3.

**Figure S7.**
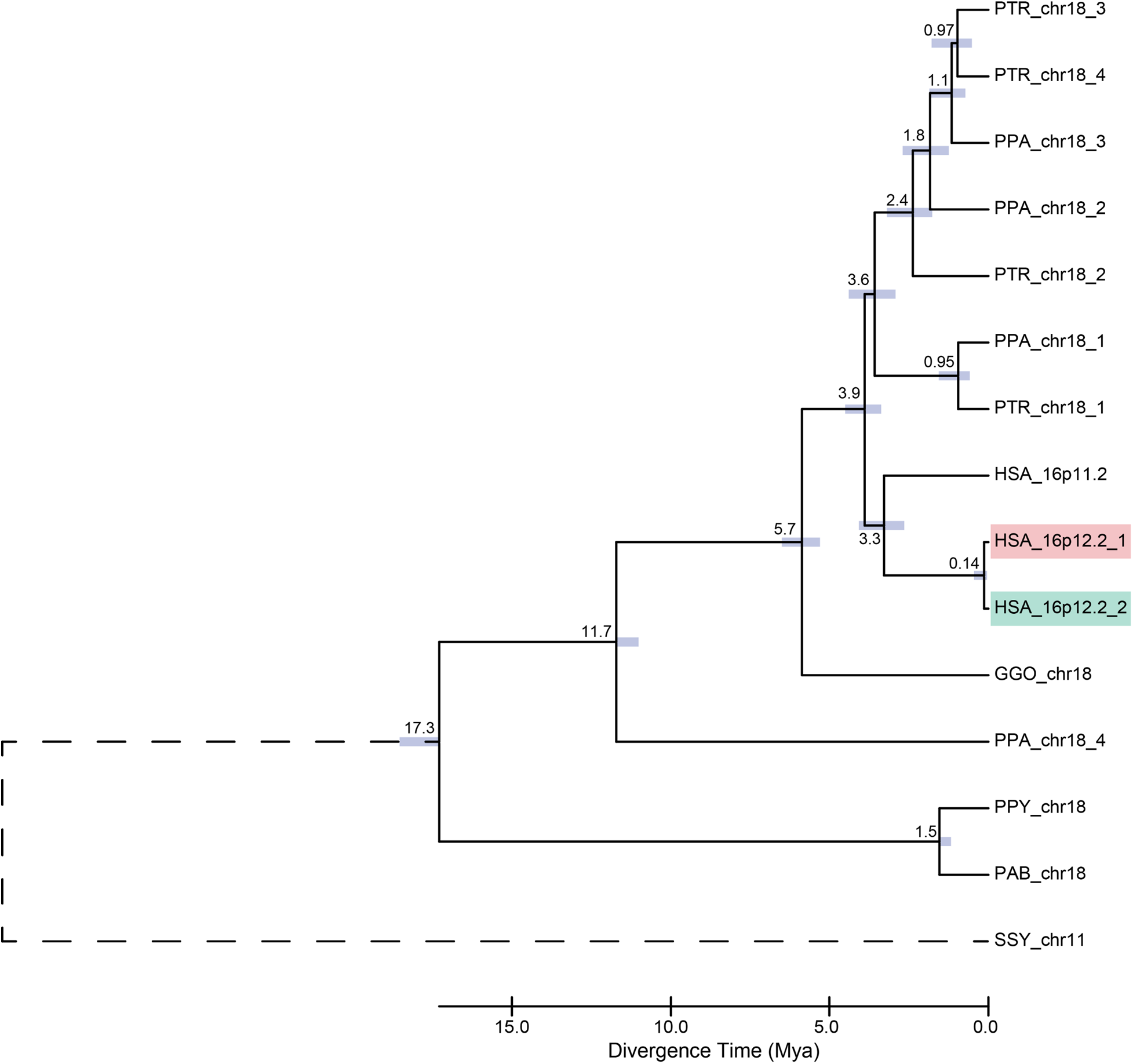
Phylogenetic tree for duplicon D8. Phylogenetic tree for D8. Dotted line indicates outgroup and colored bars indicate 95% confidence interval. Human 16p12.2 copies are shaded by the SD blocks they reside in: pink, BP1; blue, BP2; teal, BP3.

**Figure S8.**
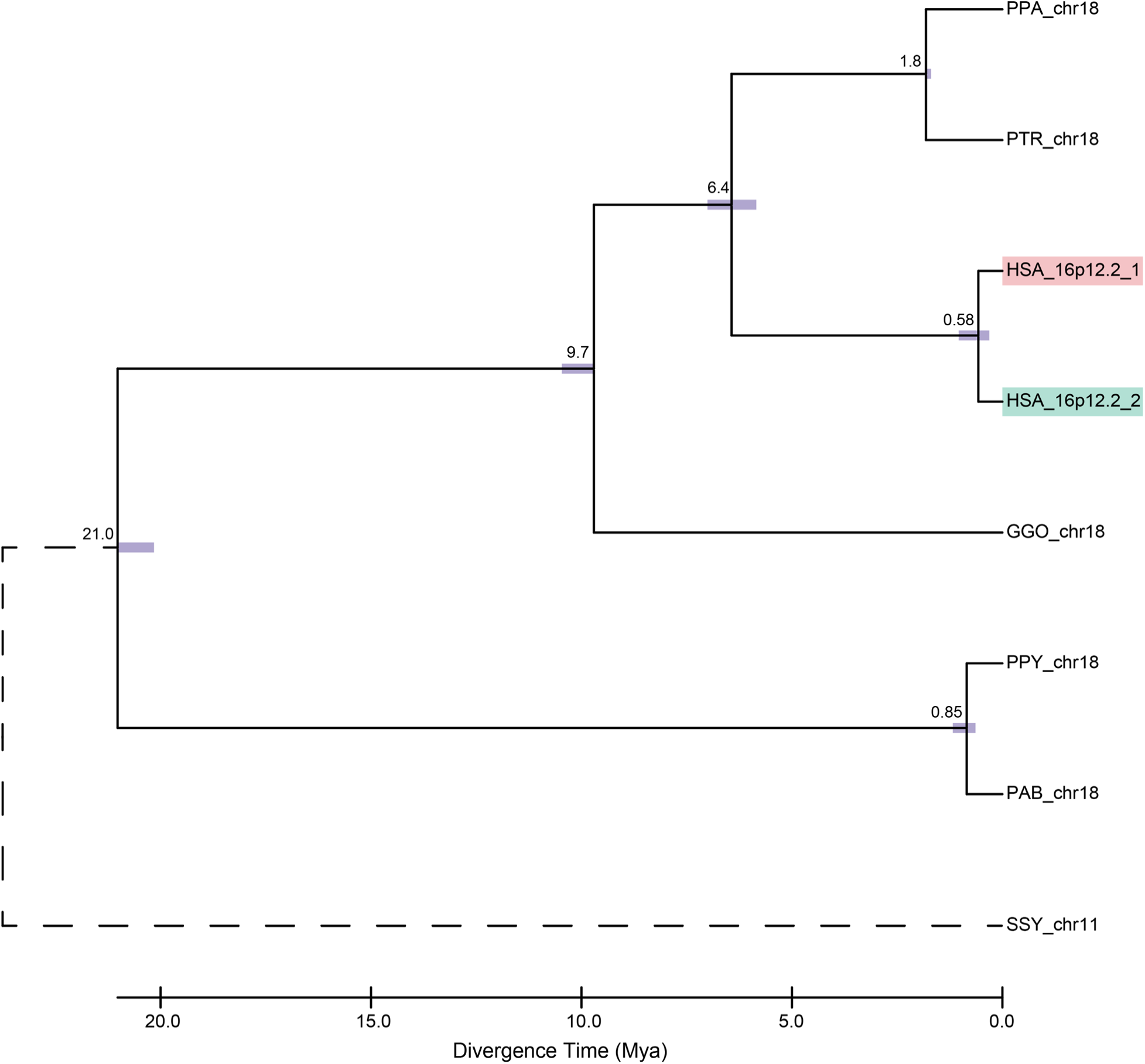
Phylogenetic tree for duplicon D9. Phylogenetic tree for D9. Dotted line indicates outgroup and colored bars indicate 95% confidence interval. Human 16p12.2 copies are shaded by the SD blocks they reside in: pink, BP1; blue, BP2; teal, BP3.

**Figure S9.**
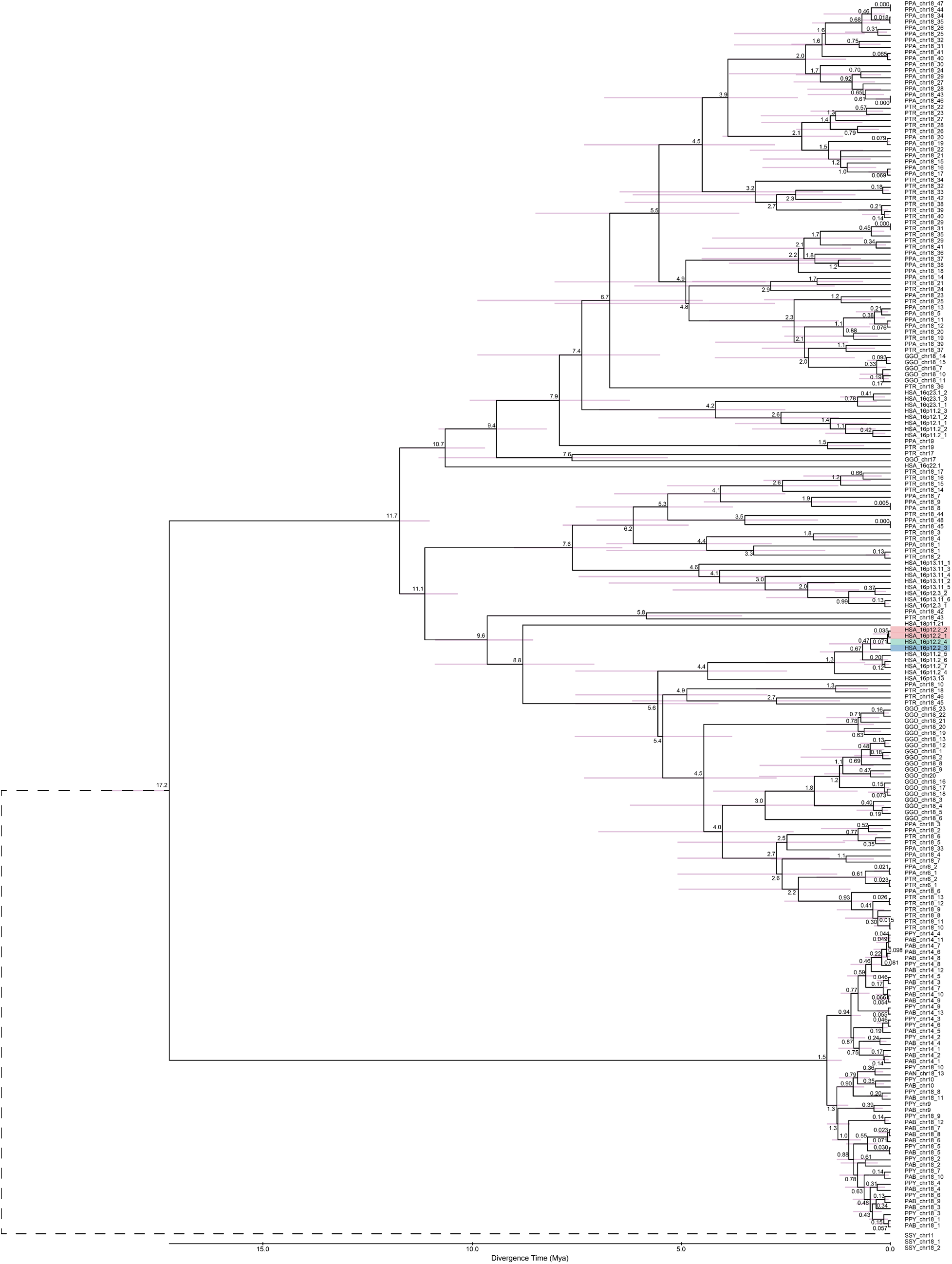
Phylogenetic tree for duplicon D10. Phylogenetic tree for D10. Dotted line indicates outgroup and colored bars indicate 95% confidence interval. Human 16p12.2 copies are shaded by the SD blocks they reside in: pink, BP1; blue, BP2; teal, BP3.

**Figure S10.**
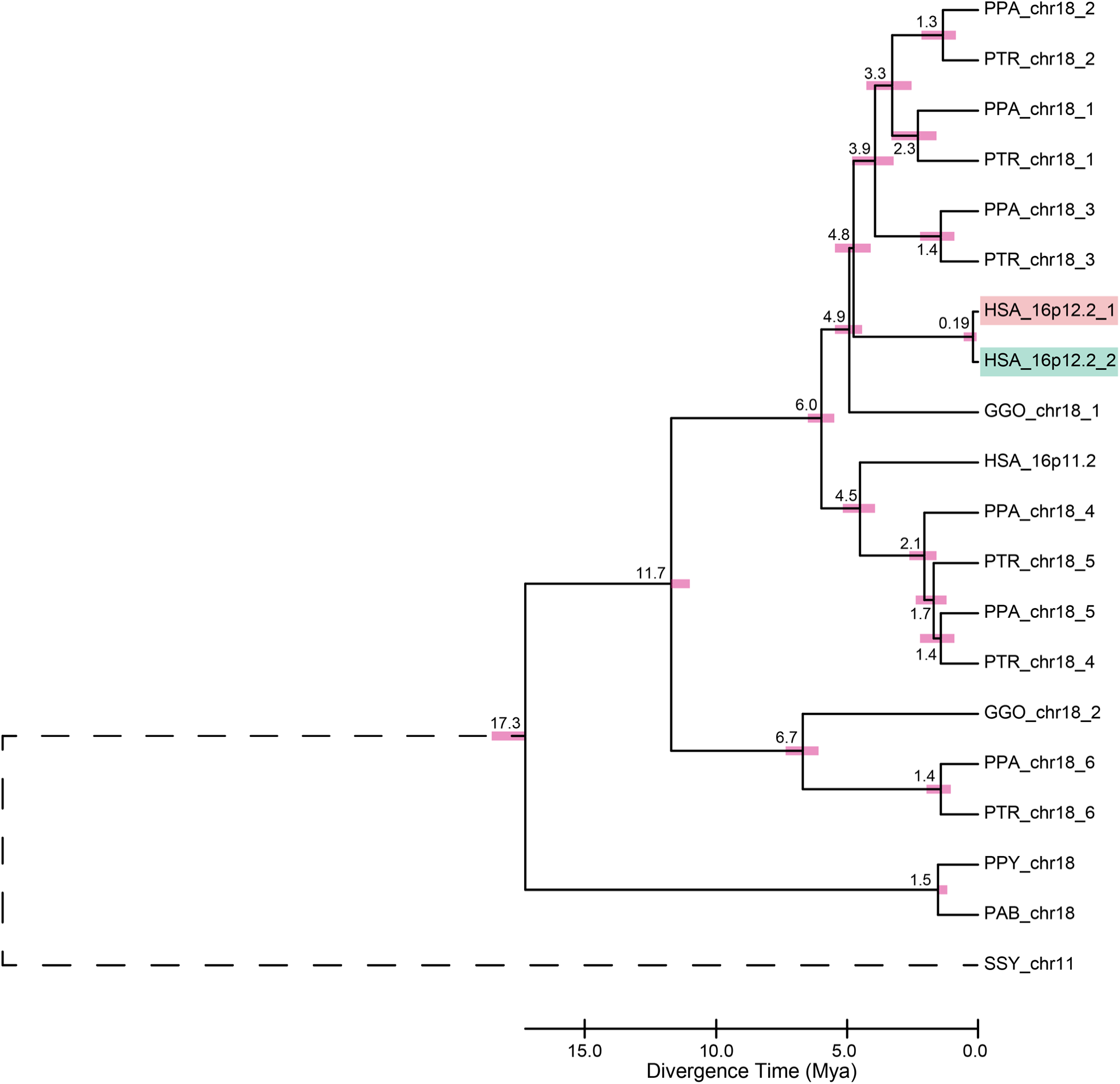
Phylogenetic tree for duplicon D11. Phylogenetic tree for D11. Dotted line indicates outgroup and colored bars indicate 95% confidence interval. Human 16p12.2 copies are shaded by the SD blocks they reside in: pink, BP1; blue, BP2; teal, BP3.

## ONLINE METHODS

### Patient Recruitment

We performed phenotyping and long read sequencing on 19 individuals from seven families with the 16p12.1 deletion, consisting of six trios and one singleton proband. All families were identified through prior clinical genetic testing for genetic causes of developmental disorders in probands. Informed consent was obtained according to a protocol approved by the Pennsylvania State University Institutional Review Board (IRB #STUDY00000278).

### Phenotyping

For probands, we used standardized questionnaires to assess parent- or guardian-reported phenotypes and grouped these phenotypes inro broad phenotypic domains (**Table S15**). Developmental delays were determined by assessing the attainment of specific developmental milestones (**Table S16**). Probands with at least one milestone achieved after the expected age were determined to have developmental delays. For parents, phenotypes were assessed using interpreted responses to a standardized questionnaire. Participants were determined to have a phenotype if they responded “Yes” to at least one question associated with a given phenotype (**Table S17**).

### Long read sequencing

Whole blood samples from all subjects were sent to Psomagen (Rockville, MD, USA) for DNA extraction, library preparation, and long read sequencing. High molecular weight (HMW) DNA extraction was performed using the Quick-DNA HMW MagBead Kit (Zymo Research, Irvine, CA, USA). A PacBio Revio DNA HiFi (Menlo Park, CA, USA) library was generated for SMRT (single molecule, real-time) sequencing using the PacBio Revio system. Samples were sequenced with an average of 5.157 M reads/sample and an average read length of 17,762 bases/read.

### Genome assembly

We used hifiasm v0.16.0^48^ with default parameters to generate haplotype-phased assemblies for each subject. For probands, we used yak v0.1-r69-dirty^67^ to count parental k-mers and supplied these k-mers to hifiasm to generate parent-phased assemblies.

### Long read mapping

We aligned long reads from 16p12.1 deletion families to the CM13 reference genome using Winnowmap2^68^ v2.03. We first used meryl to identify k-mers (k=15) in CHM13 and then used Winnowmap2 with parameter *-ax map-pb* to align long reads to CHM13.

### HPRC and HGSVC genomes

Assembled genomes from HPRC^43^ were downloaded from the HPRC data explorer (https://data.humanpangenome.org/assemblies). Assembled genomes from HGSVC^44^ were downloaded from the International Genome Sample Resource (https://www.internationalgenome.org/data-portal/data-collection/hgsvc2).

### Extracting the 16p12.2 locus

We followed the protocol described by Porubsky et al.^36^ to isolate the 16p12.2 locus. Briefly, we used minimap2 v2.30^69,70^ with parameters *-x asm20 -secondary=no -s 25000 -c* to align assembled genomes to the CHM13-T2T reference genome. We then used rustybam v0.1.34 (https://github.com/mrvollger/rustybam) to extract a region from the paf file that aligned to chr16:21000000-23500000. We removed any samples where the alignment spanned multiple contigs or multiple non-contiguous sequences within a single contig or the start or end of the 16p12.2 region was within 50kb of the end of the contig. If the entire locus mapped inverted to CHM13-T2T, it was assumed to be an assembly error and manually reversed.

### Identifying duplicated sequence with DupMasker

To identify the amount of duplicated sequence in each haplotype, we first masked high copy repeats in each 16p12.2 locus using RepeatMasker v4.1.9^71^ with parameters *-s -xsmall -e ncbi - species human*. We then used DupMasker v4.1.9^72^ with *-engine ncbi* to identify segmental duplications. We parsed the DupMasker output and compared the annotated duplicated length to the length of the locus in each sample to determine the amount of duplicated and unique sequence in each haplotype.

### Identifying SD blocks

To identify SD blocks in CHM13-T2T, we downloaded previous SEDEF SD calls for CHM13-T2T^6^ from the UCSC Genome Browser (https://genome.ucsc.edu)^73^. We then collapsed SD blocks separated by < 100 bp. This resulted in three SD blocks: chr16: 21277817-21797731 (BP1), chr16: 22280072-22487895 (BP2), and chr16: 22633819-22977140 (BP3).

To identify SD blocks in each haplotype, we followed the protocol from Porubsky et al.^36^. After extracting the 16p12.2 locus in each sample, we aligned the locus to itself using minimap2 v2.30 with parameters *-DP -k 19 -w 19 -m 200 -c --eqx*. We then used SVByEye^74^ in R v4.4.3 to filter the resultant PAF file using *filterPaf* with parameters *min.mapq=0, is.selfaln=T, min.align.len=10000, min.selfaln.dist=20000*. We then collapsed gaps < 50 kbp to identify ranges of SD blocks in each haplotype.

To determine SD block identity in each haplotype, we used three complementary methods. *First*, we aligned the haplotype SD blocks to the SD blocks from CHM13-T2T using minimap 2 v2.30 with parameters *-x map-hifi -a*. We then filtered for high-quality alignments with samtools^75^ *view* with parameters *-F 2308 -q 20*. We then assigned SD block identity based on these alignments. If the high-quality alignments were not sufficient to assign identity, we also used the unfiltered alignments. *Second,* we used MASH^76^ *triangle* to calculate the Jaccard distance between the haplotype SD blocks and the CHM13-T2T SD blocks and assigned identity based on the smallest distance. *Third*, we examined the relative positions of the SD blocks to one another based on those in CHM13. We finally compared the assignments from all three methods. Assignments agreed upon by at least two of the three methods were used for downstream analysis. For 16 samples, no two methods agreed on assignments. These samples were manually reviewed, and identities were assigned for 15 samples, while one was dropped from downstream analysis of SD block identity.

### Identifying SD block clusters

After extracting the SD loci from each sample, we used PBBG v0.7.4^77^ to build a separate pangenome graphs of all haplotypes for each SD block with parameters *-p 99.9 -s 1k*. We then used the ODGI v0.9.2-0-gbe6a0202^78^ command *similarity* with the *-d* parameter to get the Jaccard distance between all pairs of haplotypes for each SD block. We then used scipy v1.16.3^79^ in Python v3.13.9 to cluster haplotypes on the Jaccard distance using *scipy.cluster.hierarchy.hc.average*. Resulting dendrograms were manually inspected to determine the cut-points to assign cluster identity.

### Identifying duplicons

To identify duplicons, we used PGGB v0.7.4 with the same parameters as above to create a single pangenome graph for all SD block haplotypes in human samples (each SD block in all samples was made into a separate path in the graph). We then used ODGI v0.9.2-0-gbe6a0202 *untangle* with parameters *-m 20000 -n 10 -j 0.85* to identify units of at least 20kb of shared sequence between haplotypes. The output of this command was parsed to identify 11 duplicon units that appeared multiple times in CHM13-T2T (**Table S2**).

We then extracted the FASTA sequences of all duplicons copies identified in CHM13-T2T and generated a multiple-sequence alignment for each duplicon using MAFFT v7.526^80^ with the *–auto* parameter and identified the longest stretch with total gaps < 500 bp. We note that any duplicons mapped in the reverse direction were reverse complemented before alignment. We then used EMBOSS v6.6.0.0^81^ to identify the consensus sequence for each duplicon. This consensus sequence was then mapped back to all haplotypes using minimap2 with parameters *- cx asm20 -p 0.5 –eqx*. Only mappings where at least 95% of the nucleotides matched were kept for downstream analyses (**Table S14**).

### Inversion detection

We used PAV v2.4.6^44^ to identify structural variants in all 570 human haplotypes using the 16p12.2 locus from CHM13 as a reference. PAV was run with parameter *--nt* using singularity v1.4.1 and the PAV v2.4.6 image available from DockerHub (https://hub.docker.com/r/becklab/pav/tags). Inversion breakpoints on CHM13 were mapped to the identified SD blocks in CHM13 (see *Identifying SD blocks*) and identified duplicons in CHM13 (see *Identifying duplicons*) to categorize inversions.

### Variant calling in 16p12.1 deletion families

We used PAV v2.4.6^44^ to identify variants in the assembled 16p12.1 deletion family genomes, including those that failed the initial QC criteria, using the same parameters described above for inversion detection using the entire CHM13 genome as a reference. PAV correctly called the deletion in nine of the 10 deletion samples, and we used the breakpoints reported by PAV as the putative breakpoints for the deletion in these nine samples.

### Identifying recombination events

To identify recombination events, we first identified heterozygous variants called by PAV in probands in the 16p12.2 region. We then used LRPhase v1.1.2^82^ and reads aligned to CHM13 to identify parental-informative reads from the 16p12.1 deletion haplotype in probands. We then isolated heterozygous variants in the parent of origin for the deletion allele for each proband called by PAV and used LRPhase to assign each read from the proband to a haplotype in the parent of origin. We then manually examined the pattern of reads in the 16p12.2 region to identify potential recombination events.

### Identifying high copy repeats around 16p12.1 deletion breakpoints

We used high copy repeats identified by Hoyt et al.^51^ in CHM13 to examine the field of high copy repeats around the deletion breakpoints. We found the nearest repeats(s) around the PAV-reported deletion breakpoints for each family and plotted those repeats against the breakpoints reported by PAV.

### Identifying local sequence around 16p12.1 deletion breakpoints

To compare the sequences around the deletion breakpoints in all samples, we extracted the deletion breakpoints on CHM13 and the breakpoint location on assembled genomes from the PAV deletion calls for each sample. We used bedtools2 v2.31.1^83^ to extract 50 bp up and downstream of the breakpoint location in each sample genome and the first and last 50 bp of the deletion region in CHM13.

### Primate genomes

We analyzed the 16p12.2 locus in recent releases of six ape genomes from Yoo et al.^45^. Genomes were downloaded from NCBI with the following assembly IDs: GCF_028858775.2 (Chimpanzee), GCF_029289425.2 (Bonobo), GCF_029281585.2 (Gorilla), GCF_028885625.2 (Bornean Orangutan), GCF_028885655.2 (Sumatran Orangutan), and GCF_028878055.3 (Siamang). We additionally downloaded a recent rhesus macaque genome (GCF_049350105.2) from Zhang et al^46^.

### Comparative 16p12.2 locus architecture in primates

To compare 16p12.2 locus architecture in humans and other primates, we extracted the 16p12.2 locus from the primate genomes using the same method described for the human haplotypes (see *Extracting the 16p12.2 locus*), with minor adjustments. For primate genomes, we took the largest contiguous sequence aligned to the locus with gap <500 kbp. Due to the translocation in the orangutan genomes, an additional 500 kbp was included downstream of the contiguous sequence in these species. We then used PGGB v0.7.4 with parameters *-I 95 -s 1k* to create a pangenome graph of the primate loci and ODGI v0.9.2-0-gbe6a0202 *untangle* with parameters *-m 1000 -n 1* to identify orthologous regions in the primate loci relative to CHM13-T2T. Region architecture was additionally examined by manual inspection of genes in the region.

### Segmental duplications and duplications in primate genomes

We identified segmental duplications in primate genomes using the same process as used for the human haplotypes (see *Identifying SD blocks*). We identified duplicated sequence in primate genomes using the same method as for the human haplotypes (see *Identifying duplicated sequence with DupMasker*), while updating the DupMasker *species* flag for each species. We note that the 500 kbp downstream of the contiguous sequence in orangutan was excluded from calculations of unique and duplicated sequence length.

### Mapping duplicons in primates

We used the same steps described for human haplotypes (see *Identifying duplicons*) to align duplicon consensus sequences to primate genomes. For primate genomes, we used a more lenient nucleotide match requirement than for human haplotypes. Only alignments where 50% of the nucleotides matched the consensus sequence were kept for downstream analysis (**Table S8**, **S12-13**). As a result of this, there are two duplicons (D2 and D7) with more copies in the 16p12.2 locus in this analysis (**Table S8**) than in previous analyses of human haplotypes (**Table S3**).

### Constructing duplicon phylogenies

From the orthologous duplicon copies identified in *Mapping duplicons in primates*, we created a multiple sequence alignment for all duplicon copies identified across species using MAFFT MAFFT v7.526^80^ with the *–auto* parameter. Any duplicons mapped in the reverse direction were reverse complemented before alignment. We then used trimAl v1.5.rev1^84^ to remove noisy sequences and IQ-TREE v3.0.1^85–87^ to generate phylogenetic trees with parameters *-T AUTO -m MFP -alrt 1000 -bb 1000 –runs 5*. We additionally provided either the macaque, siamang, or orangutan copies as outgroups for analysis, depending on which species had copies of each duplicon (**Fig. S2-10**). We then used MEGA v12.1.2^88^ to calculate divergence times on the trees using a Hasegawa-Kishino-Yano model with four gamma-distributed categories and all sites. Divergence times were calibrated on species divergence times estimated by Yoo et al. (**Table S18**).

## Notes

### Competing Interest Statement

The authors have declared no competing interest.

